# Different RNA recognition by ProQ and FinO depends on the sequence surrounding intrinsic terminator hairpins

**DOI:** 10.1101/2024.07.24.604972

**Authors:** Maria D. Mamońska, Maciej M. Basczok, Ewa M. Stein, Mikołaj Olejniczak

## Abstract

*Escherichia coli* ProQ and FinO proteins both have RNA-binding FinO domains, which bind to intrinsic transcription terminators, but each protein recognizes distinct RNAs. To explore how ProQ and FinO discriminate between RNAs we transplanted sequences surrounding terminator hairpins between RNAs specific for each protein, and compared their binding to ProQ, the isolated FinO domain of ProQ (ProQ^NTD^), and FinO. The results showed that the binding specificity of chimeric RNAs towards ProQ, ProQ^NTD^, or FinO was determined by the origin of the transplanted sequence. Further analysis showed that the sequence surrounding the terminator hairpin, including a purine-purine mismatch, in natural RNA ligands of FinO and in chimeric RNAs weakened their binding by ProQ^NTD^. Overall, our studies suggest that the discrimination between RNAs by ProQ and FinO is determined by RNA sequence elements surrounding the intrinsic terminator hairpin.

## INTRODUCTION

RNA binding proteins play important roles in gene expression regulation dependent on small RNAs (sRNAs) in bacteria (Wagner and Romby 2015; Gorski et al. 2017; Holmqvist and Vogel 2018; Hor et al. 2020; Aoyama and Storz 2023). A well-studied example is the matchmaker protein Hfq, which promotes interactions between sRNAs and mRNAs, rearranges RNA structure, and affects RNA stability (Soper and Woodson 2008; Schu et al. 2015; Updegrove et al. 2016; Andrade et al. 2018; Kavita et al. 2018; Roca et al. 2022; Malecka and Woodson 2024). Another example are FinO-domain proteins, which are present alongside Hfq in numerous β- and γ-proteobacteria, including species important for human health (Glover et al. 2015; Attaiech et al. 2017; Olejniczak and Storz 2017; Holmqvist et al. 2020). The FinO-domain proteins are involved in diverse physiological processes, such as plasmid conjugation (Glover et al. 2015), natural transformation (Attaiech et al. 2016), osmoregulation (Kerr et al. 2014; Melamed et al. 2020), adaptation to available nutrients (El Mouali et al. 2021b; Katz et al. 2021), flagellar assembly (Rizvanovic et al. 2021), persister cells formation (Rizvanovic et al. 2022), and virulence (Westermann et al. 2019; Bergman et al. 2024). However, the molecular mechanisms of the contributions of the FinO-domain proteins to these processes are not fully understood.

The FinO-domain proteins consist of the core FinO domain and additional N- or C-terminal extensions (Glover et al. 2015; Attaiech et al. 2017; Olejniczak and Storz 2017; Holmqvist et al. 2020). Despite limited sequence identity the FinO domains from different proteins have similar overall structure consisting of five α-helical segments (Ghetu et al. 2000; Chaulk et al. 2010; Gonzalez et al. 2017; Immer et al. 2020; Kim et al. 2022). Several studies showed that a FinO domain is the part of these proteins that specifically recognizes those RNA molecules, which contain intrinsic transcription terminators (Ghetu et al. 2002; Chaulk et al. 2010; Chaulk et al. 2011; Attaiech et al. 2016; Gonzalez et al. 2017; Bauriedl et al. 2020; Pandey et al. 2020; Stein et al. 2020; Kim et al. 2022). On the other hand, the C-terminal extensions of *Legionella pneumophila* RocC and *Salmonella enterica* ProQ are important for their physiological functions (Attaiech et al. 2016; El Mouali et al. 2021b; Rizvanovic et al. 2021), and likely also contribute to non-specific RNA binding (Gonzalez et al. 2017; Stein et al. 2020).

The intrinsic transcription terminators constitute the binding sites of FinO-domain proteins in their RNA ligands. The GRAD-seq study, which identified several hundred RNAs bound by ProQ in *S. enterica*, showed that ProQ recognizes structured motifs in bound RNAs (Smirnov et al. 2016). Further global profiling studies of RNA binding by *S. enterica* and *E. coli* ProQ using CLIP-seq (Holmqvist et al. 2018), by *E. coli* ProQ using RIL-seq (Melamed et al. 2020), and by *Neisseria meningitidis* minimal ProQ (NMB1681) using CLIP-seq (Bauriedl et al. 2020) showed that typical RNA binding sites of ProQ were GC-rich and followed by uridine-rich sequences, which is consistent with intrinsic transcription terminators. The studies using purified components also showed that F-like plasmid FinO protein, *L. pneumophila* RocC and *E. coli* ProQ bound tightly to intrinsic terminator hairpins with adjacent single-stranded regions, which are derived from their native RNA ligands (Jerome and Frost 1999; Attaiech et al. 2016; Stein et al. 2023). Additionally, it was observed that the RNAs bound by ProQ contained an A-rich sequence motif on the 5ʹ side of the terminator that prevented their binding by Hfq, which is another global RNA binding protein in *E. coli* (Stein et al. 2020). Further studies showed more detailed picture of the recognition of the terminator structures by FinO domain proteins. The recent X-ray study showed that the FinO domain of *L. pneumophila* RocC protein recognized the lower part of the stem of the terminator hairpin, and its single-stranded 3ʹ-terminal tail (Kim et al. 2022). It was observed that the F-like plasmid FinO protein also recognized the lower part of the terminator of FinP RNA (Arthur et al. 2011) and the 3ʹ tail (Jerome and Frost 1999). Additionally, the contribution of the stem of the terminator hairpin to RNA binding by *S. enterica* and *E. coli* ProQ was shown using mutations disrupting or shortening this region (Holmqvist et al. 2018; Stein et al. 2020), and the contribution of the sequences surrounding the terminator was shown using truncation experiments (Stein et al. 2020; Stein et al. 2023).

The RNA binding site is located on the concave face of the FinO domain (Ghetu et al. 2002; Pandey et al. 2020; Kim et al. 2022; Stein et al. 2023). The recent X-ray crystallography study showed that the terminator hairpin of RocR RNA is bound on the concave face of the FinO domain of *L. pneumophila* RocC protein (Kim et al. 2022). In this site several residues of the N-terminal part of α-helix 5 contact the double-stranded base of the terminator hairpin, while other conserved residues form hydrogen bonds with terminal nucleotides of the 3ʹ polypyrimidine tail (Kim et al. 2022). The RNA interactions with the FinO domain were first shown directly using the crosslinking of the F-like plasmid FinO protein binding to a fragment of FinP RNA, which revealed contacts including arginine and lysine residues on the concave face of the FinO domain (Ghetu et al. 2002). The involvement of the FinO domain of *E. coli* ProQ in RNA binding was also supported by hydrogen deuterium exchange studies (Gonzalez et al. 2017). Additionally, the concave face of the FinO domain of *E. coli* ProQ was indicated as the RNA binding site by analyzing the binding of the ProQ mutants in bacterial cells using bacterial three-hybrid assay (Pandey et al. 2020), and *in vitro* using gelshift assay (Stein et al. 2023). Substitutions of several amino acids on the concave face of the FinO domains of *S. enteric*a ProQ and *L. pneumophila* RocC were also identified by mutagenesis studies exploring the physiological outcomes of the mutations, which confirms their essential importance for the function of FinO-domain proteins (Attaiech et al. 2016; El Mouali et al. 2021b; Rizvanovic et al. 2021).

The FinO-domain proteins recognize specific sets of RNAs in bacterial cells (Attaiech et al. 2016; Smirnov et al. 2016; Holmqvist et al. 2018; Bauriedl et al. 2020; Gerovac et al. 2020; Melamed et al. 2020; El Mouali et al. 2021a). Some FinO-domain proteins bind hundreds of RNAs (Smirnov et al. 2016; Bauriedl et al. 2020; Holmqvist et al. 2020; Melamed et al. 2020), while others bind few RNAs (Attaiech et al. 2016; Gerovac et al. 2020; El Mouali et al. 2021a). Interestingly, *E. coli* and *S. enterica* have both a global RNA binding protein ProQ, which binds numerous RNAs, including regulatory RNAs and mRNAs (Smirnov et al. 2016; Holmqvist et al. 2018; Melamed et al. 2020), and a narrow-specificity RNA binding protein FinO, which binds only two regulatory RNAs, FinP and RepX (El Mouali et al. 2021a). The chromosomally encoded ProQ protein is composed of the N-terminal FinO domain, a positively charged linker, and the C-terminal Tudor domain (Smith et al. 2004; Smith et al. 2007; Chaulk et al. 2011; Gonzalez et al. 2017). The F-like plasmid encoded FinO protein consists of the FinO-domain accompanied by a positively charged N-terminal extension (Ghetu et al. 2000). Because both ProQ and FinO recognize intrinsic transcription terminators as their main binding motifs in RNAs (Jerome and Frost 1999; Arthur et al. 2011; Holmqvist et al. 2018; Melamed et al. 2020; Stein et al. 2020), it is not clear how their specificity of RNA recognition is determined.

To find out how ProQ and FinO recognize their respective RNA ligands we transplanted sequence motifs surrounding intrinsic terminator hairpins between RNAs specific for each protein and measured how it affected their binding affinities to each protein. The results of these experiments provided new insights on the role of RNA sequences adjacent to intrinsic terminators in enabling the discrimination between RNAs by ProQ and FinO.

## RESULTS

### Comparison of the 3ʹ terminal sequences of top RNA ligands of ProQ and FinO

To better understand how ProQ and FinO distinguish between preferred RNAs, we analyzed the sequences and structures of top RNA ligands of each protein (Suppl. Figs S1-S3), which were previously identified using global profiling methods (Holmqvist et al. 2018; Melamed et al. 2020; El Mouali et al. 2021a). We used *RNAStructure* software to compare the sequences and secondary structures of this region for two sets of top 20 RNAs bound by ProQ, which were identified by CLIP-seq or RIL-seq in *E. coli* (Holmqvist et al. 2018; Melamed et al. 2020), as well as for *E. coli* FinP and *S. enterica* RepX, which are the ligands of F-like plasmid FinO protein (van Biesen and Frost 1994; Jerome and Frost 1999; El Mouali et al. 2021a) (Suppl. Figs S1-S3). Because previous studies showed that FinO domains bind RNAs at the site consisting of the lower part of the intrinsic terminator hairpin and surrounding sequence (Arthur et al. 2011; Stein et al. 2020; Kim et al. 2022), we predicted the structures of RNA fragments consisting of the terminator hairpin, the 10-nt upstream region, and the 3ʹ tail (Suppl. Figs S1-S3). In this analysis the lower end of the terminator hairpin was defined as the closing G-C or C-G base pair of the hairpin, which was directly adjacent to the 3ʹ tail consisting mainly of U residues.

At first we compared the sequences of the three lowest base pairs of the terminator hairpins in the RNA ligands of ProQ and FinO. Out of the 33 unique RNAs identified by RIL-seq or CLIP-seq as top 20 ligands of ProQ (Holmqvist et al. 2018; Melamed et al. 2020), 24 RNAs had only G-C or C-G base pairs in all three lowest base pairs of the terminator hairpin. In 8 RNAs, two G-C or C-G pairs were present alongside one base pair containing uridine (A-U, U-A, G-U, or U-G), while in 1 RNA there was one C-G, one A-U, and one U-A base pair (Suppl. Figs. S1, S2). When the sequences of the two RNAs bound by FinO were analyzed (El Mouali et al. 2021a), we observed that in RepX RNA this region consisted of 3 G-C or C-G pairs, while in FinP it consisted of one G-C, one C-G, and one G-U pair (Suppl. Fig. S3). Hence, similar sequences are present in this part of the hairpin in RNA ligands of ProQ and FinO, which suggests that this region is not essential to discriminate between RNAs by these proteins.

Next, we compared the secondary structures formed by the 3ʹ tails and sequences upstream of the terminator hairpins. We found that in 28 of the 33 unique RNA ligands of ProQ at least two nucleotides of the 3ʹ tail closest to the base of the hairpin were involved in base-pairing with the opposing nucleotides upstream of the hairpin, while in only 5 RNAs these regions were not base paired (Suppl. Figs. S1, S2). Of the 28 RNAs with partly base-paired 3ʹ tail, 19 had nucleotides of 3ʹ tails that formed 2, 3, or 4 base pairs, which were continuous with base-pairing of the terminator stem, while in 9 others the base pairing was longer. In contrast, neither of the RNA ligands specific to FinO showed base-pairing between the nucleotides of the 3ʹ tail and the opposing nucleotides upstream of the hairpin (Suppl. Fig. S3). The fact that the majority of ProQ-specific RNAs had at least 2 nucleotides of continued base paring below the closing base pair of the terminator hairpin, while both RNAs specific for FinO did not have base pairs below the terminator hairpin, suggests that differences in this region could be related to distinct recognition of RNAs by ProQ and FinO.

Finally, we analyzed the nucleotide composition at the two positions of the 3ʹ tail and the two positions of the upstream sequence, which were closest to the base of the terminator hairpin. Among those ProQ-specific RNAs, in which these nucleotides were base-paired, most often A-U base pairs, and less often U-A or G-U base pairs were present, while in those RNAs, in which these nucleotides were unpaired, only pyrimidine-pyrimidine mismatches, C-U, U-U, or C-C were present in these two positions (Suppl. Figs. S1, S2). In contrast, in FinO-specific FinP and RepX RNAs, the closing base pair of the hairpin was neighbored by a purine-purine apposition, the A-G mismatch, followed by either C-A or U-C mismatch (Suppl. Fig. S3). In summary, in the two positions directly below the closing base pair of the terminator hairpin the top RNAs bound by ProQ have either canonical A-U or G-U base pairs or pyrimidine-pyrimidine mismatches, while purine-purine appositions are not found in these positions. On the other hand, both RNA ligands of FinO contain a purine-purine mismatch at the first position below the terminator hairpin. The fact that the sequences immediately adjacent to the closing base pair of the terminator hairpin are different between RNA ligands of ProQ and FinO in a way which could affect local RNA structure, suggests that these sequence elements could contribute to differential RNA recognition by these two proteins.

### *malM*-3ʹ RNA is specifically recognized by the ProQ protein, and FinP RNA is specifically recognized by the FinO protein

To test if ProQ and FinO distinctly bind their natural RNA ligands, we used a gelshift assay to compare the binding of each purified protein to two RNAs that were previously identified using global profiling as their natural ligands in *E. coli* cells (Fig. 1A) (Holmqvist et al. 2018; Melamed et al. 2020). One of these RNAs was the 3ʹ-UTR of *malM* mRNA (*malM*-3ʹ), which is the top RNA bound by *E. coli* ProQ identified using RIL-seq method (Melamed et al. 2020), and one of the top ligands of ProQ identified using CLIP-seq method (Holmqvist et al. 2018) (Suppl. Figs. S1, S2). The other of these RNAs was FinP RNA, which is the main RNA bound by FinO protein in *E. coli* and *S. enterica* (van Biesen and Frost 1994; El Mouali et al. 2021a) (Suppl. Fig. S3). We compared the binding of these RNAs to the full-length ProQ and to the isolated FinO domain of ProQ (ProQ^NTD^), because it was previously shown that the isolated FinO domain of ProQ specifically recognizes intrinsic transcription terminators (Chaulk et al. 2011; Pandey et al. 2020; Stein et al. 2020).

**Figure 1.**
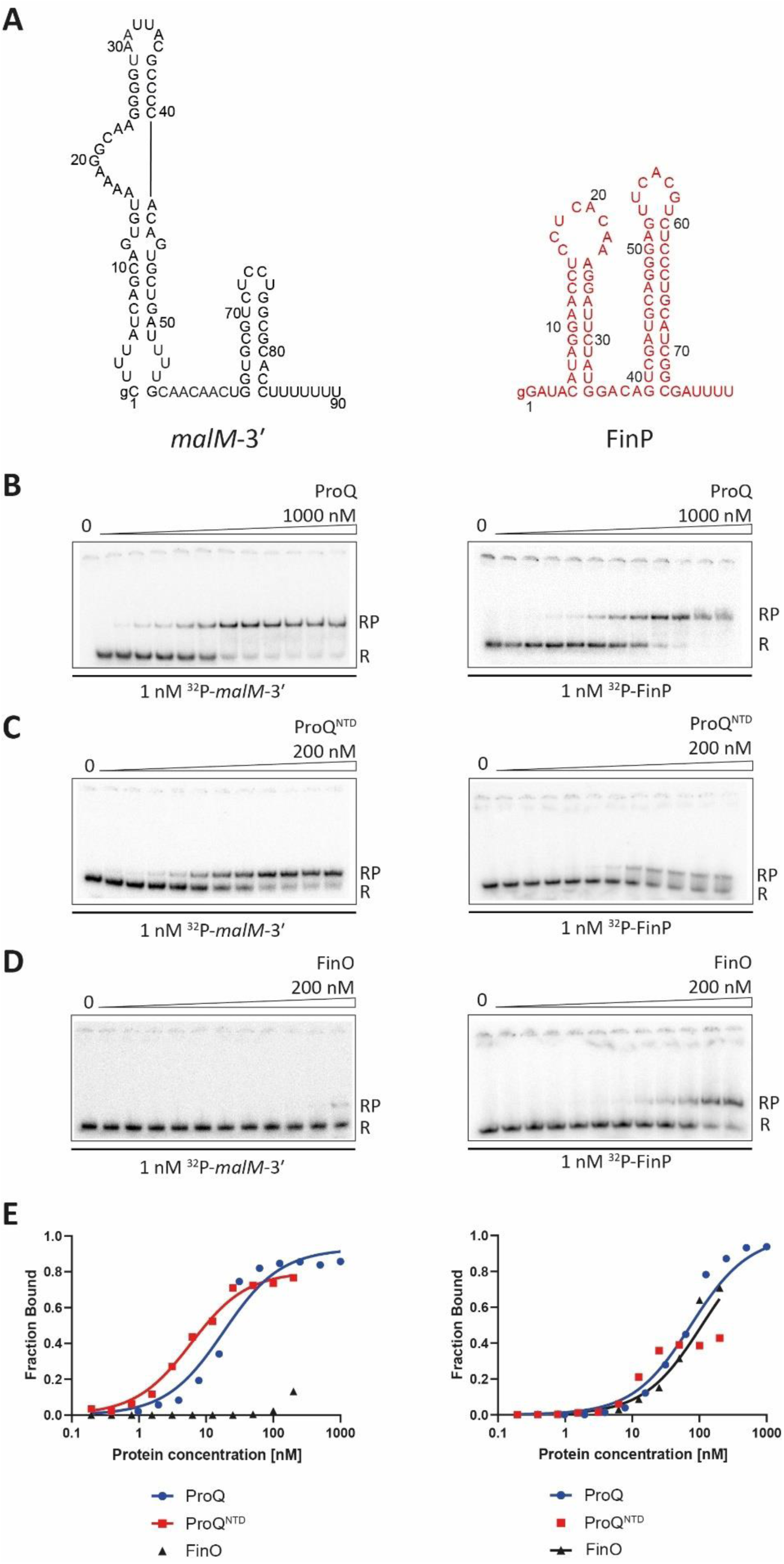
Comparison of *malM*-3’ and FinP RNA binding to ProQ, the ProQ^NTD^, and FinO. (A) Secondary structures of *malM*-3’ (black font) and FinP RNAs (red font), which were predicted using *RNAstructure* software (Reuter and Mathews 2010). The lower case g denotes guanosine residue added on 5’ ends of RNA molecules to enable T7 RNA polymerase transcription. (B) – (D) The gelshift analysis of *malM*-3’ and FinP binding to full-length ProQ (B), ProQ^NTD^ (C), and FinO (D). Free ^32^P-labeled RNA is marked as R and RNA-protein complexes as RP. (E) The plots of fraction bound data versus protein concentration from (B) – (D) are shown. The fitting of *malM*-3’ binding data using the quadratic equation provided *K*_d_ value of 18 nM for binding to ProQ, and 5.5 nM for binding to ProQ^NTD^, while the *K*_d_ value for binding to FinO was estimated as higher than 200 nM. The fitting of FinP binding data using the quadratic equation provided *K*_d_ value of 74 nM for binding to ProQ, 18 nM for binding to ProQ^NTD^, and 109 nM for binding to FinO. The average equilibrium dissociation constant (*K*_d_) values calculated from at least three independent experiments are shown in Table 1.

**Table 1.** Equilibrium RNA binding to full-length ProQ, the isolated FinO domain of ProQ (ProQ^NTD^), and FinO.

| <sup>32</sup>P-RNA | *K<sub>d</sub>* (nM) (max. fraction bound %) |  |  |
| --- | --- | --- | --- |
|  | ProQ | ProQ<sup>NTD</sup> | FinO |
| *malM*-3' | 17 ± 0.85 (91%) | 5.8 ± 0.70 (80%) | >200 (15%) |
| FinP | 79 ± 18 (85%) | 19 ± 6 (47%) | 110 ± 16 (76%) |
| *malM*-FinP | 78 ± 22 (91%) | >200 (23%) | 86 ± 17 (82%) |
| FinP-*malM* | 25 ± 6.7 (79%) | 5.3 ± 1.5 (62%) | 280 ± 110 (57%) |
| *malM*-A-GU<sub>6</sub> | n.m. | 1.9 ± 0.59 (56%) | 151 ± 40 (59%) |
| *malM*-A-GAU<sub>5</sub> | n.m. | >200 (21%) | 127 ± 18 (73%) |
| FinP-U-UAU<sub>4</sub> | n.m. | 6.2 ± 3.1 (78%) | 199 ± 73 (60%) |
| FinP-U-U<sub>6</sub> | n.m. | 9.4 ± 4.5 (82%) | 174 ± 45 (68%) |
| *cspE81*-3' | n.m. | 2.4 ± 0.27 (85%) | >200 (10%) |
| *cspE81*-FinP | n.m. | 4.9 ± 0.32 (61%) | 88 ± 39 (74%) |
| *cspE81*-FinP-stem | n.m. | >200 (37%) | 56 ± 16 (72%) |
| RepX | n.m. | >200 (6%) | 61 ± 26 (86%) |
| RepX-*malM* | n.m. | 3.2 ± 1.6 (62%) | 152 ± 67 (77%) |
| *malM*-RepX | n.m. | >200 (8%) | 160 ± 64 (57%) |

The *K<sub>d</sub>* values were obtained by fitting data to the quadratic equation. The average *K<sub>d</sub>* values with standard deviations were calculated from at least three independent experiments. When the fraction bound at the highest protein concentration used was lower than 40% the *K<sub>d</sub>* value was estimated as higher than 200 nM.
n.m. – not measured

At first, we compared the binding of full-length ProQ to both RNAs (Fig. 1B, E, Table 1). The data showed that *malM*-3ʹ RNA, which is a natural ligand of ProQ, bound ProQ with a *K*_d_ value of 17 nM, while FinP, which is a natural ligand of FinO, bound ProQ 4-fold weaker. Next, we compared the binding of these RNAs to ProQ^NTD^ (Fig. 1C, E, Table 1). *malM*-3ʹ bound ProQ^NTD^ with a *K*_d_ value of 5.8 nM, while FinP bound ProQ^NTD^ with a *K*_d_ value which was about 3-fold weaker. Additionally, the fraction of FinP bound to ProQ^NTD^ at saturation was below 50%, while that of *malM*-3ʹ saturated at 80%, which further supports weaker binding of FinP to ProQ^NTD^. In the next step, we compared the binding of both RNAs to FinO (Fig. 1D, E, Table 1). The fraction of *malM*-3ʹ bound to FinO protein was less than 15% at the maximum 200 nM concentration of FinO used, which allows estimating the *K*_d_ value as weaker than 200nm. On the other hand FinP bound FinO with the *K*_d_ value of 110 nM, while the binding saturated at more than 70% fraction bound, which confirms stronger binding of the FinO protein to its natural ligand FinP than to *malM*-3ʹ (Fig. 1D, E, Table 1). In summary, full length ProQ, ProQ^NTD^, and FinO each have tighter binding affinity to its respective natural RNA ligand than to the other RNA, which is consistent with their distinct recognition of these RNAs in bacterial cells (Holmqvist et al. 2018; Melamed et al. 2020; El Mouali et al. 2021a).

### Transplanting sequence elements surrounding the terminator hairpin from FinP into *malM*-3ʹ switches the preferred binding from ProQ to FinO

Because the analysis of sequences and secondary structures of RNA ligands of ProQ and FinO showed differences in the regions surrounding their terminator hairpins (Suppl. Figs. S1-S3), we designed chimeric constructs with these sequences transplanted between FinP and *malM*-3ʹ (Figs. 2, 3) to test if transplanted sequences would affect their binding to ProQ and FinO. In the natural sequence of *malM*-3ʹ the three nucleotides 5ʹ-adjacent to the terminator hairpin are ACU, and it has a 3ʹ terminal U_7_ tail (Fig. 1). The sequence elements transplanted from FinP into the *malM*-3ʹ body were the A residue 5ʹ-adjacent to the terminator hairpin, and the 3ʹ-terminal GAU_4_ sequence (Fig. 2A). The resulting *malM*-FinP chimera had the three nucleotides 5ʹ-adjacent to the terminator hairpin and the whole 3ʹ-terminal tail the same as in FinP.

**Figure 2.**
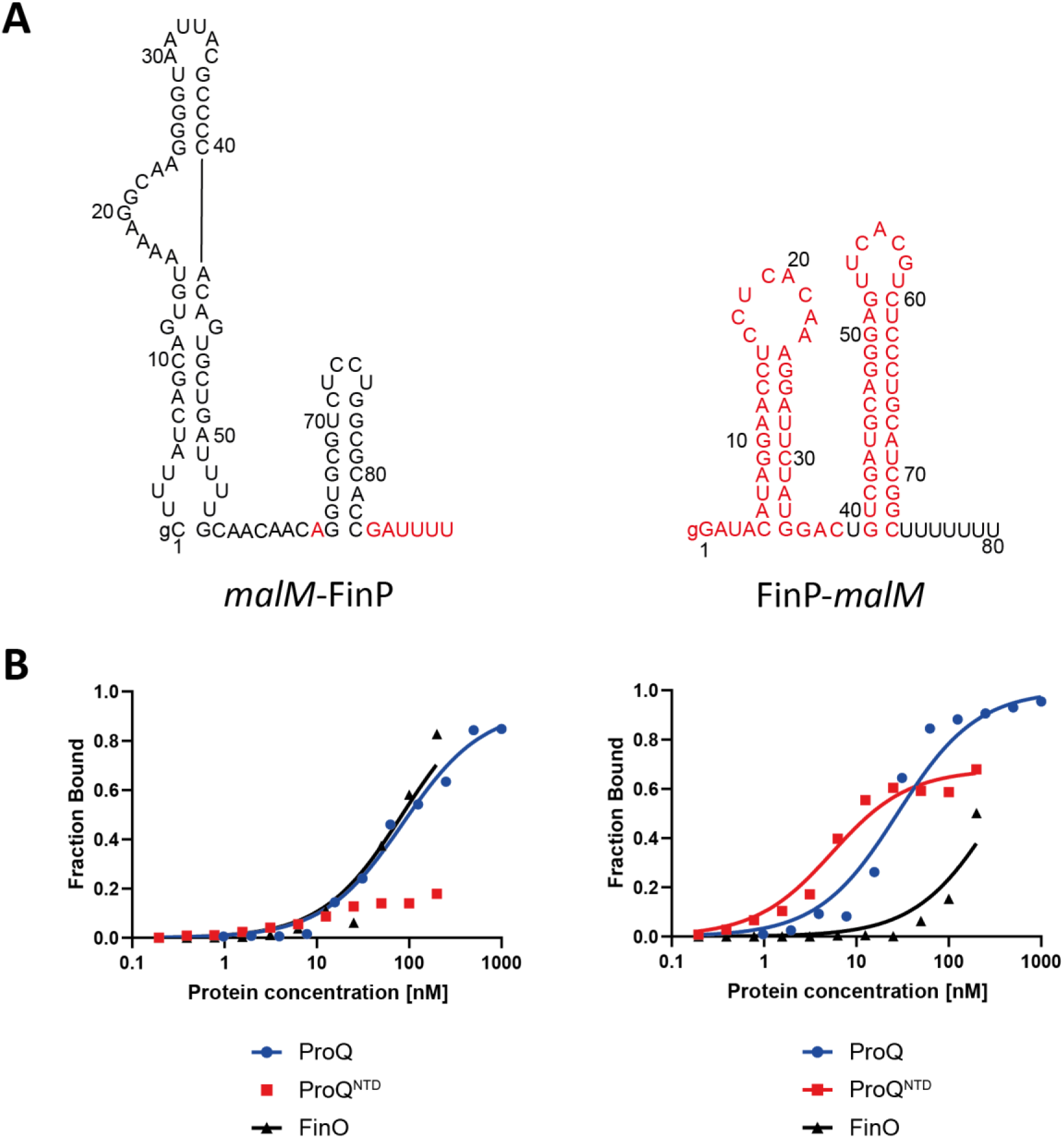
Comparison of *malM*-FinP and FinP-*malM* chimeras binding to ProQ, the ProQ^NTD^, and FinO. (A) Secondary structures of *malM*-FinP and FinP-*malM* chimeras, which were predicted using *RNAstructure* software (Reuter and Mathews 2010). The sequences originating from *malM*-3’ are shown in black font, and the sequences from FinP in red font. The lower case g denotes guanosine residue added on 5’ ends of RNA molecules to enable T7 RNA polymerase transcription. (B) The respective binding data for ProQ, the ProQ^NTD^, and FinO are shown on graphs below each RNA. The fitting of *malM*-FinP data using the quadratic equation provided *K*_d_ value of 85 nM for binding to ProQ and 83 nM for binding to FinO, while the *K*_d_ value for binding to ProQ^NTD^ was estimated as higher than 200 nM. The fitting of FinP-*malM* data using the quadratic equation provided *K*_d_ value of 27 nM for binding to ProQ, 4.9 nM for binding to ProQ^NTD^, and 327 nM for binding to FinO. Gels corresponding to the data in the plots are shown in Supplementary Figure S4. Average *K*_d_ values are shown in Table 1.

The data showed that *malM*-FinP chimera bound to full-length ProQ with similar affinity as FinP, but more than 4-fold weaker than *malM*-3ʹ (Figs. 2B, Table 1, Suppl. Fig. S4A). This detrimental effect was even stronger for the binding of *malM*-FinP chimera to ProQ^NTD^, because the fraction of *malM*-FinP bound was less than 25% at the maximum 200 nM concentration of ProQ^NTD^ used (Fig. 2B, Table 1, Suppl. Fig. S4B). Consistently, *malM*-FinP chimera bound to the FinO protein with similar affinity as FinP, and much more strongly than *malM*-3ʹ (Fig. 2B, Table 1, Suppl. Fig. S4C). In summary, these data indicated that the sequence elements of FinP transplanted into the body of *malM*-3ʹ strengthened the binding of resulting chimera to the FinO protein as compared to *malM*-3ʹ, while they weakened its binding to either ProQ or ProQ^NTD^.

### Transplanting sequence elements surrounding terminator hairpin from *malM*-3ʹ into FinP switches the preferred binding from FinO to ProQ

To test if the corresponding sequence from *malM*-3ʹ can affect RNA recognition by ProQ and FinO, we constructed a chimera, in which the sequence elements from *malM*-3ʹ were transplanted into FinP RNA to create a FinP-*malM* chimera (Fig. 2A). In the natural sequence of FinP the three nucleotides 5ʹ-adjacent to the terminator hairpin are ACA, and there is a 3ʹ terminal GAU_4_ tail (Fig. 1A). The sequence elements from *malM*-3ʹ transplanted into the FinP body were the U residue 5ʹ-adjacent to the terminator hairpin, and the 3ʹ-terminal U_7_ tail (Fig. 2A). As a result the FinP-*malM* chimera had the three nucleotides 5ʹ-adjacent to the terminator hairpin, and the whole 3ʹ-terminal tail the same as in *malM*-3ʹ.

The data showed that FinP-*malM* chimera bound ProQ similar to *malM*-3ʹ, and 3-fold tighter than FinP (Fig. 2B, Table 1, Suppl. Fig. S4A). Consistently, the FinP-*malM* chimera bound ProQ^NTD^ with the same affinity as *malM*-3ʹ, and 3-fold stronger than FinP (Fig. 2B, Table 1, Suppl. Fig. S4B). On the other hand, FinP-*malM* bound the FinO protein 2-fold more weakly than FinP, but markedly stronger than *malM*-3ʹ (Fig. 2B, Table 1, Suppl. Fig. S4C). The fact that FinP-*malM* bound FinO stronger than *malM*-3ʹ could suggest that the structure context into which the sequence elements of *malM*-3ʹ were transplanted also affects the binding of FinO. Overall, these data showed that the sequence elements adjacent to the terminator hairpin in *malM*-3ʹ direct RNA recognition by the ProQ protein, even when they are placed in the context of an RNA that is not naturally bound by ProQ.

### Dissection of *malM*-3ʹ sequence elements, which determine the binding specificity towards ProQ

To dissect which specific sequence elements within the transplanted sequence of *malM*-3ʹ are responsible for differential recognition of this RNA by ProQ and FinO, we designed two *malM*-3ʹ mutants, in which only the nucleotides closest to the hairpin were substituted for such nucleotides as are present in corresponding positions in FinP (Fig. 3A). The first of these mutants, named *malM*-A-GU_6_, had each of the nucleotides directly adjacent to the terminator hairpin on its 5ʹ and 3ʹ side substituted for opposing A and G residues, respectively. The second mutant, named *malM*-A-GAU_5_, additionally had a uridine in the second position on the 3ʹ side of the hairpin substituted for adenosine. Hence the *malM*-A-GU_6_ mutant had an A-G apposition below the terminator hairpin the same as in FinP, while the *malM*-A-GAU_5_ mutant additionally had a C-A apposition in the second position below the hairpin the same as in FinP (Figs. 1, 3A). The data showed that introducing the A-G apposition into the *malM*-3’-A-GU_6_ mutant did not markedly affect its binding to ProQ^NTD^, which had a low nanomolar affinity, although the binding saturated at below 60%, as compared to 80% observed for unmodified *malM*-3ʹ (Fig. 3B, Table 1, Suppl. Fig. S5A). On the other hand, introducing the A-G apposition very strongly improved the binding of FinO, because the affinity of the *malM*-3’-A-GU_6_ mutant to FinO was similar as that of FinP RNA (Fig. 3B, Table 1, Suppl. Fig. S5B). The additional substitution introducing the A residue in the second position on the 3ʹ side of the terminator hairpin had a strong detrimental effect on the binding of the resulting *malM*-A-GAU_5_ mutant to ProQ^NTD^ (Fig. 3B, Suppl. Fig. S5A). This effect was not caused by the shortening of the 3ʹ terminal oligoU sequence by the purine substitutions, because when the oligoU tail was extended to 7 nucleotides as in wt *malM*-3ʹ the binding of the resulting *malM*-A-GAU_7_ mutant to ProQ^NTD^ was not restored (Suppl. Fig. S6). On the other hand, the binding of *malM*-3’-A-GAU_5_ to FinO was not further improved as compared to *malM*-3’-A-GU_6_ (Fig. 3B, Suppl. Fig. S5B). Hence, in the context of *malM*-3’ body both the A-G apposition in the first position and the additional adenosine substitution leading to the C-A apposition in the second position below the terminator hairpin were necessary to abrogate the binding of ProQ^NTD^, while the A-G apposition in the first position alone was sufficient to rescue the binding to FinO.

**Figure 3.**
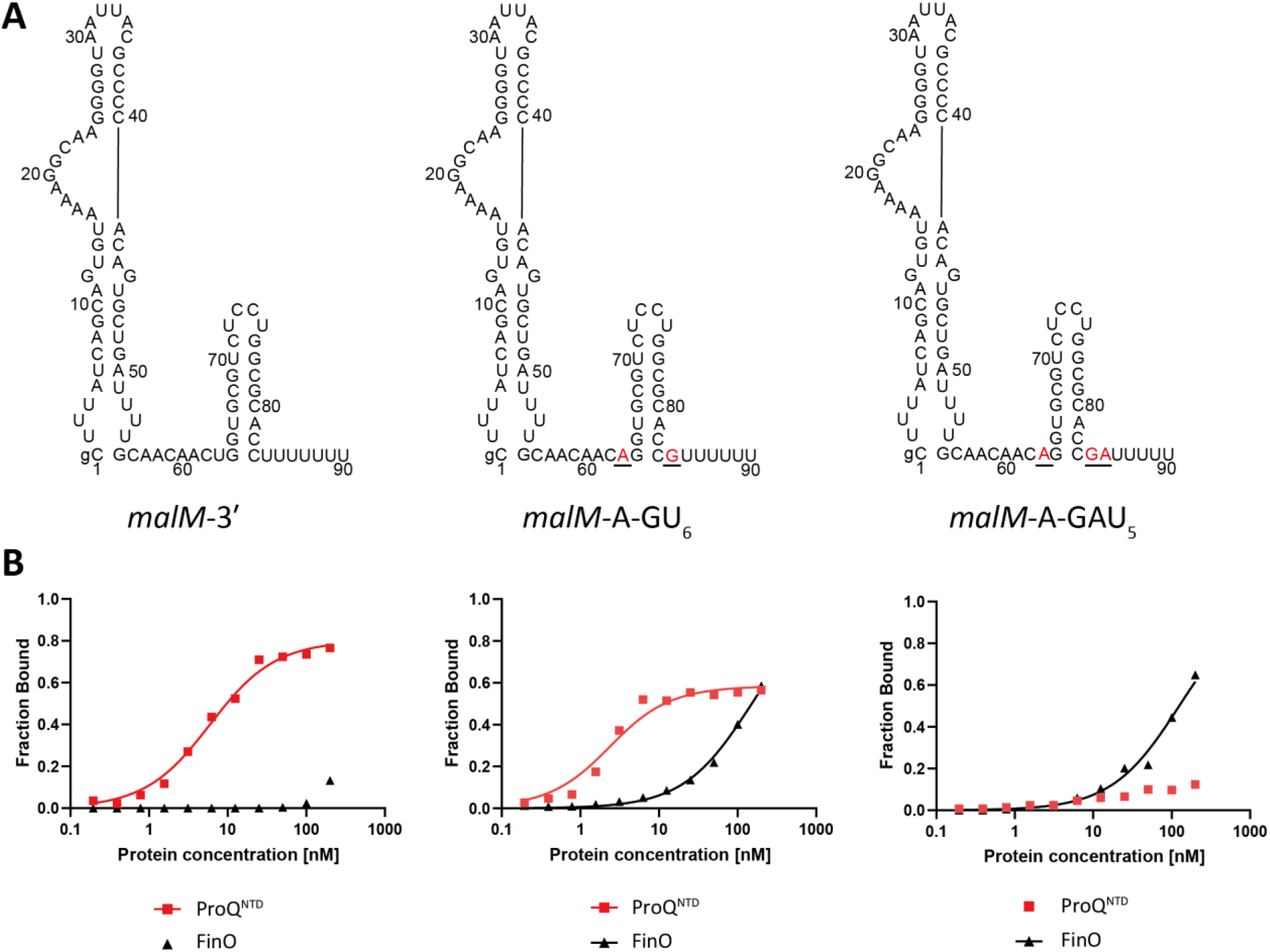
Comparison of *malM*-3’, *malM*-3’-A-GU_6_, and *malM*-3’-A-GAU_5_ binding to ProQ^NTD^, and to FinO. (A) Secondary structures of *malM*-3’, *malM*-3’-A-GU_6_, and *malM*-3’-A-GAU_5_, which were predicted using *RNAstructure* software (Reuter and Mathews 2010). The nucleotides from FinP which were substituted into *malM*-3’ are shown in red underlined font. The lower case g denotes guanosine residue added on 5’ ends of RNA molecules to enable T7 RNA polymerase transcription. (B) The respective binding data for ProQ^NTD^ and FinO are shown on graphs below each RNA. The fitting of *malM*-3’-A-GU_6_ data using the quadratic equation provided *K*_d_ value of 1.8 nM for binding to ProQ^NTD^, and 151 nM for binding to FinO. The fitting of *malM*-3’-A-GAU_5_ data using the quadratic equation provided *K*_d_ value of 122 nM for binding to FinO, while the *K*_d_ value for binding to ProQ^NTD^ was estimated as higher than 200 nM. The data shown for *malM*-3’ are the same as in Figure 1. Gels corresponding to the data in the plots are shown in Supplementary Figure S5. Average *K*_d_ values are shown in Table 1.

### Dissection of FinP sequence elements, which determine the binding specificity towards FinO

To explore whether the sequence elements at the two positions closest to the terminator hairpin of FinP are also involved in differential recognition of this RNA by ProQ and FinO, we designed two mutants of FinP (Fig. 4, Table 1). The first mutant, named FinP-U-UAU_4_, only had both purines in the first positions below the terminator hairpin substituted with uridines, which are present in the corresponding locations in *malM*-3ʹ (Fig. 4A). The second mutant, named FinP-U-U_6_, additionally had the adenosine in the second-position below the hairpin on the 3ʹ side replaced with a uridine, which is present in the corresponding position in *malM*-3ʹ (Fig. 4 A). The data showed that the substitution of the A-G apposition to the U-U apposition in the FinP-U-UAU_4_ mutant was sufficient to very strongly improve its binding by ProQ^NTD^ with the *K*_d_ value in a low nanomolar range (Fig. 4B, Table 1, Suppl. Fig. S7A). On the other hand, these mutations had only a weak approximately 2-fold detrimental effect on the binding of this RNA to FinO (Fig. 4B, Table 1, Suppl. Fig. S7B). The additional substitution of adenosine to uridine in the second position on the 3ʹ side of the hairpin in FinP-U-U_6_ did not further affect the binding to either protein beyond the changes already caused by substitution of the two purines in the FinP-U-UAU_4_ mutant (Fig. 4B, Table 1, Suppl. Fig. S7). To test whether the tight binding of FinP-U-U_6_ to ProQ^NTD^ was caused by the displacement of the purine nucleotides or by the resulting lengthening of the 3ʹ terminal oligoU sequence we compared its binding with that of FinP-U-U_4_ mutant which had the number of 3ʹ terminal uridine residues the same as in FinP (Suppl. Fig. S8). The data showed that shortening the 3ʹ U tail to 4 residues in FinP-U-U_4_ mutant did not weaken the binding of ProQ^NTD^, which suggests that the main reason for the improved binding of the FinP-U-U_6_ mutant to ProQ^NTD^ is the absence of the purine-purine apposition neighboring the terminator hairpin, rather than the lengthening of the stretch of uridine residues at the 3ʹ end. Additionally, the data showed markedly weaker binding of the FinP-U-U_4_ mutant to FinO, which suggests that the binding of the FinO protein is more negatively affected than the binding of ProQ^NTD^ by the shortening of the total 3ʹ tail length in the context of the FinP body (Suppl. Fig. S8). In summary, these data suggest that the presence of the A-G apposition directly below the closing G-C base pair of FinP RNA terminator hairpin is a negative determinant preventing its binding to ProQ.

**Figure 4.**
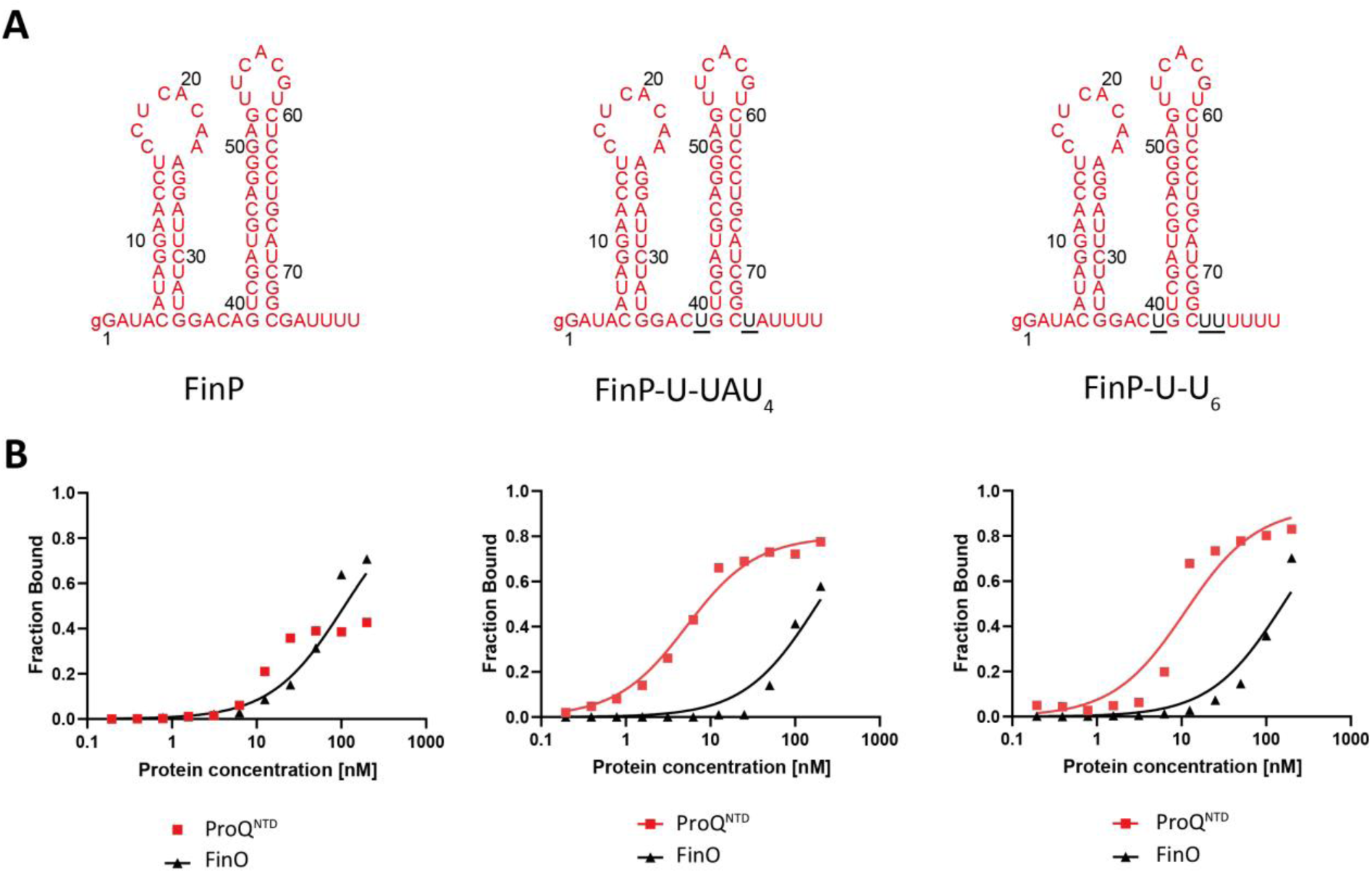
Comparison of FinP, FinP-U-UAU_4_, and FinP-U-U_6_ binding to ProQ^NTD^, and to FinO. (A) Secondary structures of FinP, FinP-U-UAU_4_, and FinP-U-U_6_, which were predicted using *RNAstructure* software (Reuter and Mathews 2010). The nucleotides from *malM*-3’ which were substituted into FinP are shown in black underlined font. The lower case g denotes guanosine residue added on 5’ ends of RNA molecules to enable T7 RNA polymerase transcription. (B) The respective binding data for ProQ^NTD^ and FinO are shown on graphs below each RNA. The fitting of FinP-U-UAU_4_ data using the quadratic equation provided *K*_d_ value of 4.6 nM for binding to ProQ^NTD^, and 184 nM for binding to FinO. The fitting of FinP-U-U_6_ data using the quadratic equation provided *K*_d_ value of 11 nM for binding to ProQ^NTD^, and 160 nM for binding to FinO. The data shown for FinP are the same as in Figure 1. Gels corresponding to the data in the plots are shown in Supplementary Figure S7. Average *K*_d_ values are shown in Table 1.

### Sequence elements transplanted from FinP into the body of another ProQ ligand, *cspE*-3ʹ RNA, prevent the binding to ProQ^NTD^

The data presented above showed that transplanting sequence elements surrounding the terminator hairpin of FinP into the corresponding positions in *malM*-3ʹ body switched its binding preference from ProQ^NTD^ to FinO (Figs. 1-3, Table 1). However, *malM*-3ʹ belongs to only few RNAs among the top ligands of ProQ, which naturally contain a pyrimidine-pyrimidine mismatch immediately below the closing G-C or C-G base pair of the terminator hairpin (Suppl. Figs. S1, S2). The majority of top RNA ligands of ProQ have an A-U base pair in this position. An example is *cspE*-3ʹ, which is among RNAs bound by ProQ identified in CLIPseq and RILseq studies (Holmqvist et al. 2018; Melamed et al. 2020). To test if the binding specificity of 81-nt long *cspE*81-3ʹ is also dependent on sequences surrounding the transcription terminator we transplanted the sequences surrounding the terminator hairpin from FinP into an 81-nt *cspE*-3ʹ (Fig. 5A). In the natural sequence of *cspE*-3ʹ the four nucleotides 5ʹ-adjacent to the terminator hairpin are all adenosines, and it has a 3ʹ terminal U_8_ sequence, of which 4 uridines nearest to the closing C-G base pair of the terminator hairpin are base-paired with opposing adenosines (Fig. 2A). The sequence elements transplanted from FinP into the *cspE*81-3ʹ body were the C residue in the second position on the 5ʹ-side to the terminator hairpin, and the 3ʹ-terminal GAU_4_ sequence (Fig. 2A). As a result the *cspE*81-FinP chimera had the three nucleotides 5ʹ-adjacent to the terminator hairpin and the whole 3ʹ-terminal tail the same as in FinP.

**Figure 5.**
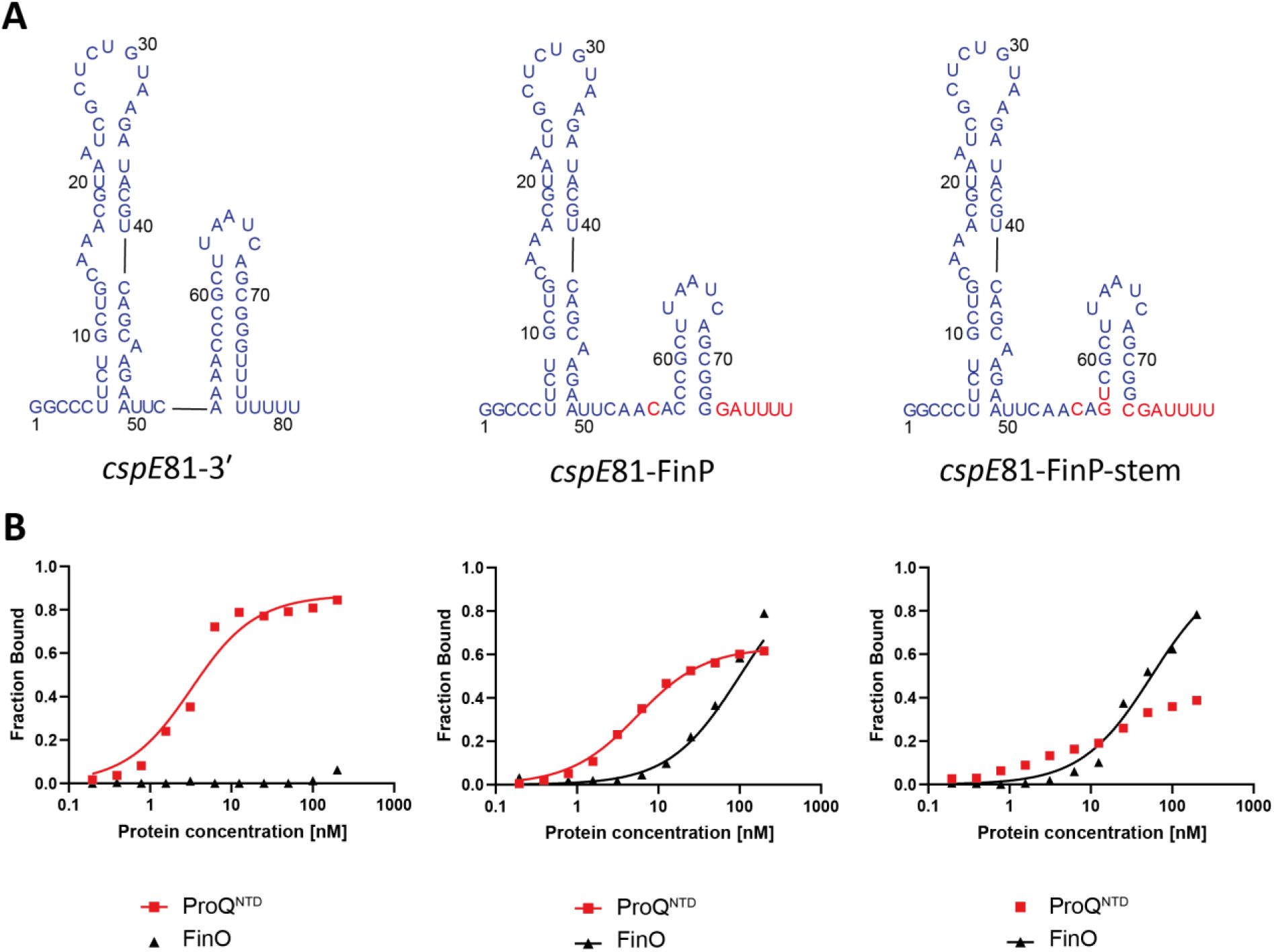
Comparison of *cspE*81-3’, *cspE*81-FinP chimera, and *cspE*81-FinP-stem chimera binding to the ProQ^NTD^, and to FinO. (A) Secondary structures of *cspE*81-3’, *cspE*81-FinP chimera, and *cspE*81-FinP-stem chimera, which were predicted using *RNAstructure* software (Reuter and Mathews 2010). The sequences originating from *cspE*81-3’ are shown in dark blue font, and the sequences from FinP in red font. The lower case g denotes guanosine residue added on 5’ ends of RNA molecules to enable T7 RNA polymerase transcription. (B) The respective binding data for ProQ^NTD^ and FinO are shown on graphs below each RNA. The fitting of *cspE*81-3’ data using the quadratic equation provided *K*_d_ value of 2.7 nM for binding to ProQ^NTD^, while the *K*_d_ value for binding to FinO was estimated as higher than 200 nM. The fitting of *cspE*81-FinP data using the quadratic equation provided *K*_d_ value of 5.0 nM for binding to ProQ^NTD^, and 99 nM for binding to FinO. The fitting of *cspE*81-FinP-stem data using the quadratic equation provided *K*_d_ value of 55 nM for binding to FinO, while the *K*_d_ value for binding to ProQ^NTD^ was estimated as higher than 200 nM. Gels corresponding to the data in the plots are shown in Supplementary Figure S9. Average *K*_d_ values are shown in Table 1.

The data showed that *cspE*81-3ʹ bound to ProQ^NTD^ with subnanomolar *K*_d_, which was even 2-fold tighter than that of *malM*-3ʹ (Fig. 5B, Table 1, Suppl. Fig. S9A). On the other hand its binding to FinO protein was negligible with maximum fraction bound of 10% at the 200 nM concentration of FinO (Fig. 5B, Table 1, Suppl. Fig. S9B). The introduction of substitutions, which made the sequence surrounding the terminator in the *cspE*81-FinP chimera the same as in FinP, resulted in moderate, 2-fold weakening of *cspE*81-FinP binding to ProQ^NTD^, and also resulted in a lower maximum fraction bound. On the other hand, these substitutions restored the binding of *cspE*81-FinP RNA to FinO to the affinity similar as that of FinP RNA. Because we noted that the lowest two base pairs of the terminator hairpin differ between *cspE*81-3ʹ and FinP, we next made additional substitutions into *cspE*81-3ʹ to ensure that in the resulting *cspE*81-3ʹ-FinP-stem chimera not only the surrounding sequence but also the two lowest base pairs of terminator hairpin are the same as in FinP (Fig. 5A). While these substitutions did not markedly affect the binding of *cspE*81-FinP-stem chimera to FinO they resulted in further weakening of its binding to ProQ^NTD^ (Fig. 5B, Table 1, Suppl. Fig. S9). In summary, these data show that in the context of the *cspE*81-3ʹ body both the sequence surrounding the terminator hairpin and the two lowest base pairs of the hairpin affect the recognition of *cspE*81-3ʹ by ProQ and FinO. This suggests that while the recognition of *malM*-3ʹ and *cspE*81-3ʹ by ProQ and FinO is dependent on the same structural features the details of the recognition differ between these two RNAs.

### Sequence elements transplanted from another FinO ligand, RepX RNA, into the body of *malM*-3ʹ prevent the binding to ProQ^NTD^

Because recent studies in *S. enterica* showed that the F-like plasmid FinO protein binds specifically not only to FinP but also to another RNA, named RepX (El Mouali et al. 2021a), we tested if sequence elements surrounding the terminator hairpin of RepX serve similar role in determining RNA recognition by FinO as we observed for FinP (Figs. 1-3, Table 1). For this we designed two chimeric constructs. In one, the sequence elements surrounding the terminator hairpin in *malM*-3ʹ were transplanted into RepX (Fig. 6A). The sequence elements transplanted into the RepX-*malM* chimera from *malM*-3ʹ were the ACU sequence on the 5ʹ side of the terminator hairpin, and the U_7_ tail on the 3ʹ side of the hairpin (Fig. 6A). In the other chimera, the sequence elements surrounding the terminator hairpin in RepX were transplanted into *malM*-3ʹ (Fig. 6A). The sequence elements transplanted into the *malM*-RepX chimera from RepX were the UUA sequence on the 5ʹ side of the terminator hairpin, and the GCUCU tail on the 3ʹ side of the hairpin (Fig. 3A). Of note, the nucleotide residues nearest to the closing base pair of the terminator hairpin of RepX, A and G, are the same as in FinP.

**Figure 6.**
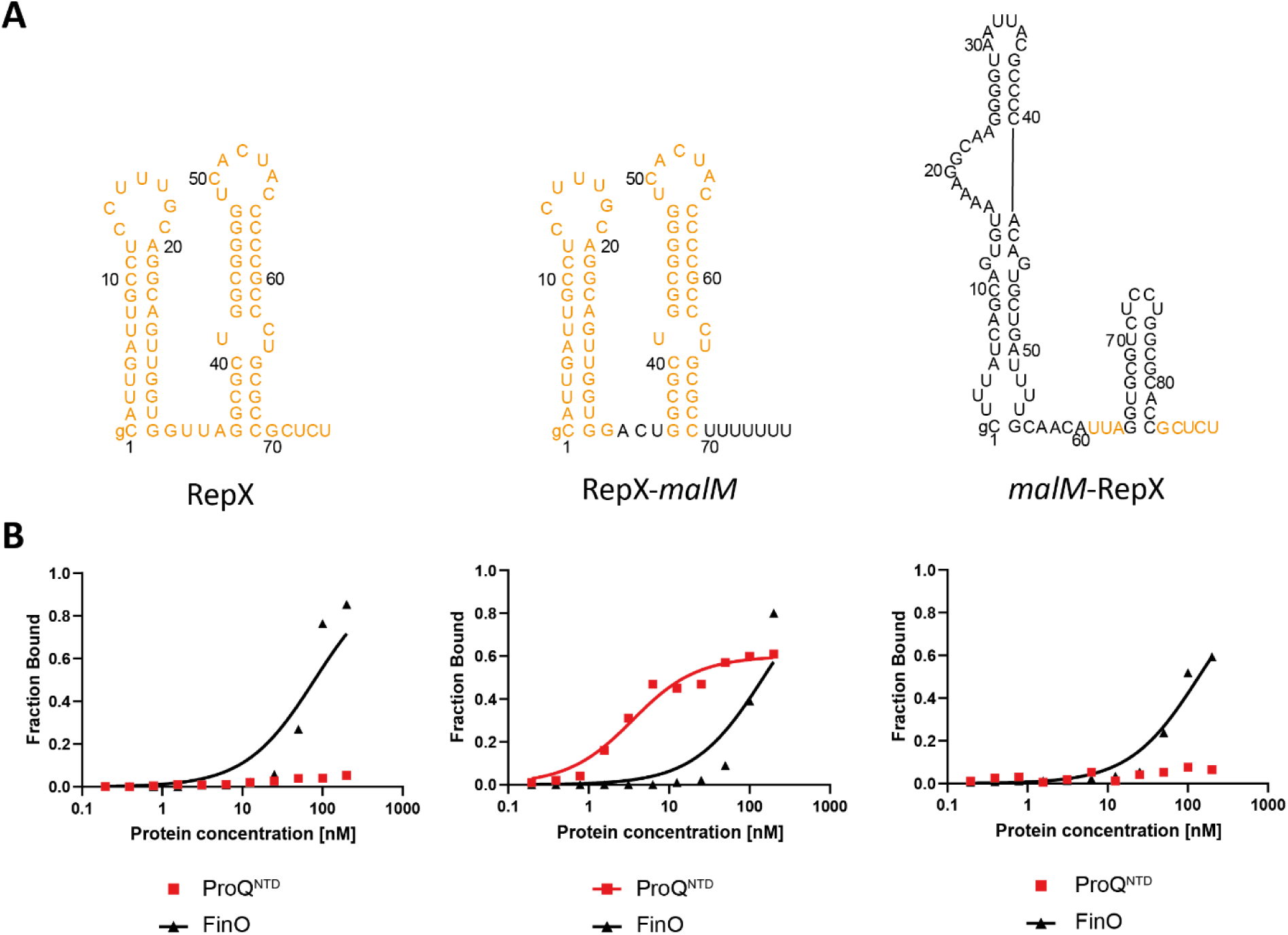
Comparison of *malM*-3’, RepX, and *malM*-RepX chimera binding to the ProQ^NTD^, and to FinO. (A) Secondary structures of *malM*-3’, RepX, and *malM*-RepX chimera, which were predicted using RNAstructure software (Reuter and Mathews 2010). The sequences originating from *malM*-3’ are shown in black font, and the sequences from RepX in orange font. The lower case g denotes guanosine residue added on 5’ ends of RNA molecules to enable T7 RNA polymerase transcription. (B) The respective binding data for ProQ^NTD^ and FinO are shown on graphs below each RNA. The fitting of RepX data using the quadratic equation provided *K*_d_ value of 79 nM for binding to FinO, while the *K*_d_ value for binding to ProQ^NTD^ was estimated as higher than 200 nM. The fitting of RepX-*malM* data using the quadratic equation provided *K*_d_ value of 3.0 nM for binding to ProQ^NTD^, and 148 nM for binding to FinO. The fitting of *malM*-RepX data using the quadratic equation provided *K*_d_ value of 137 nM for binding to FinO, while the *K*_d_ value for binding to ProQ^NTD^ was estimated as higher than 200 nM. The data shown for *malM*-3’ are the same as in Figure 1. Gels corresponding to the data in the plots are shown in Supplementary Figure S10. Average *K*_d_ values are shown in Table 1.

The data showed that the binding of wt RepX chimera to ProQ^NTD^ was almost undetectable up to 200 nM concentration of the protein (Fig. 6B, Table 1, Suppl. Fig. S10A), which is consistent with the weak binding of FinP to ProQ and ProQ^NTD^ (Figs. 1C,E, 2B, Table 1, Suppl. Fig. S4). At the same time, RepX bound the FinO protein with a *K*_d_ value of 77 nM (Fig. 6B, Table 1, Suppl. Fig. S10B), which is similar to the affinity of FinP binding to FinO (Fig. 1D, E, Table 1). In contrast, the RepX-*malM* chimera bound strongly to ProQ^NTD^ with a *K*_d_ value similar to that of *malM*-3ʹ, although its binding to FinO was not much affected by the transplantation (Fig. 6B, Table 1, Suppl. Fig. S10B). This showed that removing the RepX sequence adjacent to terminator hairpin enabled the tight binding of RepX-*malM* chimera to ProQ^NTD^. When the binding of the reverse chimera, *malM*-RepX, was measured the data showed that its binding to ProQ^NTD^ was almost undetectable, which is similar to RepX (Fig. 6B, Table 1, Suppl. Fig. S10A). On the other hand *malM*-RepX chimera bound FinO with a *K*_d_ value of 160 nM, which is only 2-fold weaker than for RepX (Fig. 6B, Table 1, Suppl. Fig. S10B). Overall, these data showed that sequence elements surrounding the intrinsic terminator hairpin in RepX have similar role as the corresponding sequence elements in FinP, because they function to weaken RNA binding by the ProQ^NTD^.

## DISCUSSION

The data presented here show that the sequences surrounding the transcription terminator hairpin contribute not only to the strength of binding of FinO-domain proteins to their RNA ligands, but also to the discrimination among RNAs by these proteins (Figs. 1-6, Table 1). The recent crystal structure of the FinO domain of *L. pneumophila* RocC protein in complex with RocR RNA showed in molecular detail how the lower part of RocR hairpin and its 3ʹ polypyrimidine tail interact with amino acid residues on the concave face of the FinO domain of RocC (Kim et al. 2022). Additionally, several *in vitro* binding studies showed the importance of the lower part of the terminator hairpin and adjacent single-stranded sequences for tight RNA binding by *E. coli* ProQ (Chaulk et al. 2011; Stein et al. 2020; Stein et al. 2023) and FinO (van Biesen and Frost 1994; Jerome and Frost 1999; Arthur et al. 2011). Here, we observed that transplanting sequences surrounding the terminator hairpins between respective RNA ligands of each protein changed the binding affinities of these RNAs to ProQ and FinO in agreement with the origin of the transplanted sequence (Figs. 1-6, Table 1). This indicates that the transplanted sequences contribute to the specificity of RNA recognition by ProQ and FinO. Hence, the data presented here expand the role of this RNA region by showing that the features of RNA sequences surrounding the terminator hairpin allow discrimination between RNAs by *E. coli* ProQ and FinO.

The sequences adjacent to the terminator hairpins of natural RNA ligands of the FinO protein, FinP and RepX, prevent the binding of these RNAs to the FinO domain of ProQ. Our data showed that when natural sequences surrounding terminators were removed from either FinP or RepX, and replaced with corresponding sequence from *malM*-3ʹ, it resulted in tighter binding of chimeric RNAs to ProQ^NTD^ (Figs. 2 and 6, Table 1). Conversely, introducing the corresponding sequence from FinP into either *malM*-3ʹ or *cspE*-3ʹ, and from RepX into *malM*-3ʹ, weakened their binding to ProQ^NTD^ (Figs. 2, 3, 6, Table 1). The unusual feature of both FinP and RepX is the presence of a purine-purine mismatch immediately below the closing G-C base pair of the terminator hairpin (Suppl. Fig. S3). On the other hand, most of the top RNA ligands of ProQ have extended A-U base-pairing below the terminator stem (Suppl. Figs. S1, S2). Although A-U base-pairing is not predicted for *malM*-3ʹ, which contains a pyrimidine-pyrimidine mismatch below the terminator, the sequence on the 5ʹ side of the terminator contains several adenosines, which suggests that less stable base-pairing in this region could still form. Indeed, a pyrimidine-pyrimidine mismatch in the first position below the terminator hairpin, which conformed to the A-type double helix geometry, was observed in the structure of *L. pneumophila* RocR RNA with RocC protein (Kim et al. 2022). This suggests that the RNA binding site of the FinO domain of ProQ can accommodate RNAs, which contain A-U base-pairs and pyrimidine-pyrimidine mismatches below the closing base pair of the terminator. We hypothesize that the reason that the sequences surrounding the terminators of FinP and RepX prevent their binding by ProQ is that the purine-purine mismatch together with neighboring sequence elements disrupt the double-helical structure of this region in a way, which is not compatible with its recognition by the FinO domain of ProQ.

The sequences surrounding terminators of FinP and RepX not only serve to prevent the binding of ProQ, but also contribute to the binding of FinO. Introducing the sequences from FinP or RepX into the context of *malM*-3ʹ or *cspE*-3ʹ strengthened the binding of these RNAs to the FinO protein (Figs. 2, 3, 6, Table 1). This suggests that the sequences naturally surrounding the terminator of FinP and RepX introduced into chimeric RNAs such features that are recognized by FinO. On the other hand, when the corresponding sequences from *malM*-3ʹ were introduced into FinP or RepX it had only a moderate detrimental effect on the binding by FinO. It was previously proposed that the RNA strands surrounding the terminator of FinP are separated in the complex of FinP with FinO (Arthur et al. 2011). We hypothesize that the natural sequence of FinP, including the purine-purine mismatch, helps to facilitate the strand separation, which is why introducing such sequence into *malM*-3ʹ or *cspE*-3ʹ strengthens the binding. On the other hand, the moderate detrimental effect of replacing the natural sequence of FinP or RepX with that from *malM*-3ʹ could suggest either that the strands originating from this RNA are sufficiently separated in the context of unnatural RNA body to enable the binding by FinO, or that other parts of the FinP or RepX structure also contribute to the RNA binding by FinO. In support of the latter hypothesis it was previously observed that the sole terminator hairpin of FinP, devoid of surrounding single stranded regions was still able to bind FinO, albeit much weaker than the intact RNA (Jerome and Frost 1999). Regardless of the detailed explanation, we hypothesize that the differences in sequences around the terminator hairpins of natural RNA ligands of ProQ and FinO lead to differences in local RNA structure, thus affecting RNA recognition by each protein.

What properties of distinct FinO domain proteins make them capable of differently recognizing the same RNAs? The X-ray structure of a complex of *L. pneumophila* RocC protein with RocR RNA showed that the double-helical stem of RocR terminator hairpin and the end of its 3ʹ tail were recognized by two distinct regions of the FinO domain of RocC (Kim et al. 2022). In this structure the two terminal nucleotides of the 3ʹ polypyrimidine tail were bound by a conserved group of residues including a tyrosine and an arginine corresponding to *E. coli* ProQ residues Y70 and R80. However, because these amino acids are the same in *E. coli* ProQ and F-plasmid FinO protein, they would not be expected to be directly responsible for different RNA recognition by ProQ and FinO. On the other hand, the lower part of double-helical stem of the RocR terminator hairpin is bound by a set of hydrogen-bond forming amino acids in the N-terminal part of α-helix 5, which was named the α-helical N-cap motif (Kim et al. 2022). The N-cap motif of RocC consists of two serines, two lysines and an arginine of α-helix 5 (S70, K71, S72, K74, and R76), which are within hydrogen-bonding distance to nonbridging oxygens of the phosphates of the 3ʹ strand of the terminator stem of RocR (Kim et al. 2022). As these amino acid side chains contact the base of the terminator hairpin, and there are differences in the sequences forming corresponding α-helices between ProQ and FinO, it is possible that this region could be responsible for differential recognition of RNA ligands by ProQ and FinO.

Because the structures of the complexes of *E. coli* ProQ and FinO with their RNA ligands are not known, we overlayed the NMR structure of the FinO domain of ProQ (Gonzalez et al. 2017) and the X-ray crystal structure of FinO (Ghetu et al. 2000) on the structure of the complex of RocC with RocR (Kim et al. 2022) to visualize the surfaces of α-helices of ProQ and FinO corresponding to RocC α-helix 5, which would likely be exposed towards bound RNA (Fig. 7). For comparison, we also modeled these interactions by overlaying the ColabFold-generated structures of the FinO domain of ProQ protein and the FinO protein on the structure of the complex of RocC with RocR (Kim et al. 2022; Mirdita et al. 2022) (Suppl. Fig. S6). The orientations of amino acids side chains towards RNA were similar in the models based on ColabFold generated structures as in models based on experimentally obtained structures (Fig. 7, Suppl. Fig. S6). The analysis of the modeled interactions showed that the N-cap motif in α-helix 3 of ProQ is quite similar to that of RocR, because it contains hydrogen-bonding residues in corresponding positions (S53, K54, R58 and R62) (Fig. 7, Suppl. Fig. S11). The importance of K54, R58 and R62 for RNA binding by *E. coli* ProQ has previously been shown by mutagenesis experiments *in vivo* and *in vitro* (Pandey et al. 2020; Stein et al. 2023). On the other hand, in the FinO protein the surface of the corresponding α-helix 4 from which hydrogen-bonding residues are directed toward modeled RNA molecules is more extended and covers the whole length of this α-helix. It includes seven hydrogen-bonding residues, which side chains are oriented towards RNA double helix (H117, K118, R121, R122, K125, and R129) (Fig. 7, Suppl. Fig. S11). The importance of R121 and K125 for RNA binding by FinO has already been shown using cysteine substitutions and crosslinking (Ghetu et al. 2002). Overall, the comparison of distributions of hydrogen-bonding amino acids on ProQ α-helix 3 and corresponding FinO α-helix 4 shows marked differences, which could form the basis of unique interactions that enable each protein to differently recognize RNA molecules.

**Figure. 7.**
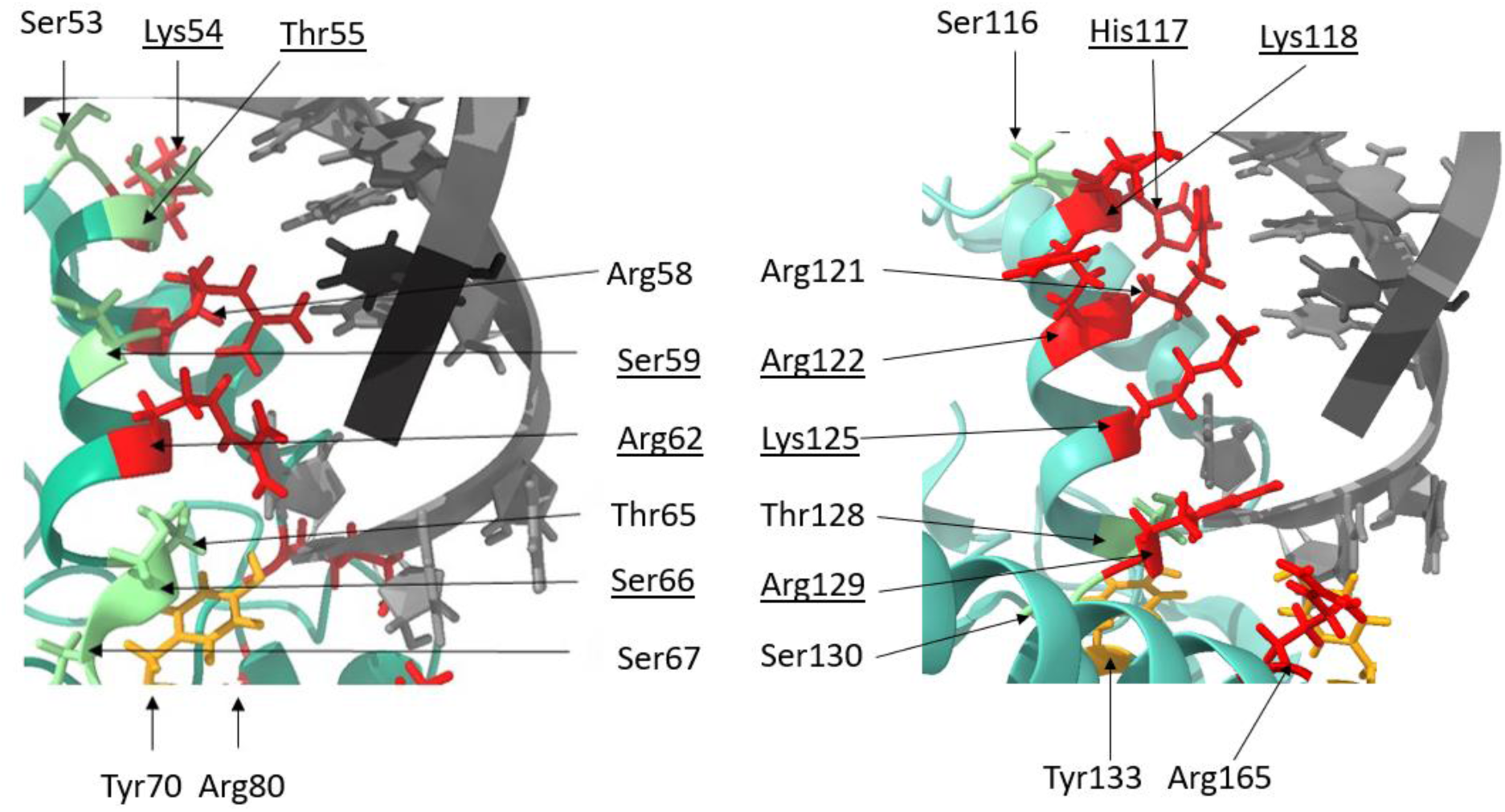
The modeling of RNA binding surfaces of *E. coli* ProQ and F-like plasmid FinO. The figure shows the α-helix 3 and surrounding region from *E. coli* ProQ (left) and the corresponding α-helix 4 from F-like plasmid FinO (right) with those amino acid residues marked, which side chains are directed towards modeled location of RNA helix. The modelling of interactions was done using Chimera X (Pettersen et al. 2021) by aligning the NMR structure of the FinO domain of *E. coli* ProQ (Gonzalez et al. 2017) and the X-ray structure of the F-like plasmid FinO protein (Ghetu et al. 2000) with the X-ray structure of the FinO domain of *L. pneumophila* RocC in complex with the terminator hairpin of RocR RNA (Kim et al. 2022). The side chains of amino acid residues located in the corresponding positions of both proteins are marked in color, with arginine, lysine and histidine residues marked in red, serine and threonine in green, and tyrosine in orange. The descriptions of corresponding amino acids are located in corresponding places on the figure. Those amino acids which are different, but located in corresponding positions, are marked by underlining. The structure of the *L. pneumophila* RocR hairpin is shown in grey.

In summary, our data suggest that the reason for the binding of distinct RNAs by ProQ and FinO in *E. coli* cells is the presence of specific sequence and structure features in the vicinity of intrinsic transcription terminators of these RNAs, which make them optimal ligands for either ProQ or FinO, while preventing the binding of the other protein. We speculate that the use of RNA features which prevent the binding of plasmid-encoded RNAs to ProQ serves to ensure that they have orthogonal access to plasmid-encoded FinO protein, and hence they do not compete with cellular RNAs for access to global RNA binding ProQ. Because it was previously observed that the A-rich sequence on the 5ʹ side of terminators of RNAs bound by ProQ weakens their binding to global RNA-binding protein Hfq, it is possible that the use of sequence elements serving as negative determinants of binding is one of mechanisms ensuring specific RNA recognition by proteins interacting with regulatory RNAs in *E. coli* and related bacteria.

## MATERIALS AND METHODS

### Preparation of RNAs

The DNA templates for *in vitro* transcription were obtained by Taq polymerase extension of chemically synthesized overlapping oligodeoxyribonucleotides (Sigma-Aldrich and Metabion) (Supplemental Table S1). RNA molecules used in this study were obtained using *in vitro* transcription with T7 RNA polymerase as described (Milligan et al. 1987; Olejniczak 2011). After transcription, RNA molecules were purified using 8 M urea polyacrylamide gel electrophoresis. RNAs were 5’-^32^P-labeled using T4 polynucleotide kinase (Thermo Scientific) and γ-^32^P ATP (Hartmann Analytic), which was followed by phenol-chloroform extraction, denaturing gel electrophoresis, and ethanol precipitation. 5’ ^32^P-labeled RNAs were stored at –20°C as 200 nM solutions.

### Expression and purification of proteins

The sequences of *Escherichia coli* ProQ protein, and its 130-aa long N-terminal domain were cloned into pET-15b vector (Novagen), and purified as previously described (Stein et al. 2020). To obtain the FinO protein the overexpression construct was prepared by cloning the coding sequence of *finO* into pET-15b vector (Novagen) using BamHI restriction site (Supplemental Table S2). The coding sequence was obtained by amplification from the template of *finO* construct in pGEX-KG vector, which was a kind gift of Prof. Mark Glover (University of Alberta). In the expression construct the coding sequence was preceded by His_6_-tag and TEV protease recognition sequence (ENLYFQ↓S). The construct was overexpressed in BL21 Δ*hfq E. coli* strain (a kind gift of Prof. Agnieszka Szalewska-Pałasz, University of Gdansk) and purified as previously described for *E. coli* ProQ (Stein et al. 2020). The molecular weight of the purified protein was determined by MALDI-TOF as 21362.1 Da, which agrees with the calculated mass of 21366.6 Da. The samples were stored in a buffer consisting of 50 mM Tris, pH 7.5, 300 mM NaCl, 10% glycerol and 1 mM EDTA, at −80°C, in 5 μl and 10 μl aliquots, and used without refreezing.

### Gelshift assays of RNA binding by ProQ, ProQ^NTD^, and FinO

RNA binding to ProQ, ProQ^NTD^ or FinO proteins was measured using electrophoretic mobility shift assays. The concentration series of ProQ, ProQ^NTD^ or FinO were made by 2-fold sequential dilutions. RNAs were denatured at 90°C for 2 min, followed by refolding on ice for 5 min. To initiate the binding reaction a ^32^P-labeled RNA (1 nM final concentration) was mixed with indicated final concentrations of ProQ, ProQ^NTD^ or FinO in the binding buffer (150 mM NaCl, 25 mM Tris-HCl pH 7.5, 5% glycerol, 1 mM MgCl_2_) and incubated for 30 min at room temperature. Incubation was performed in low-protein binding microplates, additionally pre-treated with a solution containing 0.0025% bovine serum albumin. After 30 min of incubation, 5 µl reaction aliquots were loaded onto a native 6% polyacrylamide gel (19:1), and run in 0.5× TBE at 4°C. Gels were dried using vacuum dryer, and exposed to phosphor screens, followed by data quantification using a phosphorimager and MultiGauge software (Fuji FLA-5000). In order to calculate the equilibrium dissociation constant (*K*_d_) values, the data were fit in GraphPad Prism software to the quadratic equation as described (Stein et al. 2020). Average *K*_d_ values with standard deviations were calculated from three independent experiments.

## ACKNOWLEDGEMENTS

We thank Gisela Storz for helpful discussions, and G.S., Jared Schrader, and Julia Kurzawska for critical comments on the manuscript. This work was supported by National Science Centre in Poland [grants No. 2020/39/O/NZ1/02448 and No. 2018/31/B/NZ1/02612]. Funding for open access charge: National Science Centre [2020/39/O/NZ1/02448] and Adam Mickiewicz University.

## AUTHORS CONTRIBUTIONS

M.D.M. performed all binding experiments, M.M.B. analyzed transcription terminator structures and RNA binding surfaces of ProQ and FinO, E.M.S. cloned and purified the FinO protein, M.D.M. and M.O. analyzed the data and wrote the manuscript.

