## Supplemental Materials for "Different RNA recognition by ProQ and FinO depends on the sequence surrounding intrinsic terminator hairpins"

#### SUPPLEMENTAL METHODS

##### *Prediction of secondary structures involving nucleotides adjacent to the terminator hairpins*

The secondary structures were analyzed for RNA ligands of ProQ and FinO identified in previous studies using CLIP-seq, RIL-seq and RIP-seq methods (Holmqvist et al. 2018; Melamed et al. 2020; El Mouali et al. 2021). For the analysis of RNA ligands of ProQ 20 RNA sequences with the highest averaged read counts were selected from CLIP-seq data when the peak covered the terminator loop or when it was located no further than ten nucleotides upstream of it (Suppl. Fig. S1) (Holmqvist et al. 2018). Additionally, 20 RNA sequences with the highest averaged read counts, which were annotated as 3′-UTRs or sRNAs, were selected from RILseq data obtained in LB medium (Suppl. Fig. S2) (Melamed et al. 2020). The two main RNA ligands of F-like plasmid FinO protein were analyzed, which were previously identified using RIP-seq method (Suppl. Fig. S3) (El Mouali et al. 2021). For each RNA the secondary structure was predicted for a fragment consisting of the transcription terminator hairpin together with 10 nucleotides upstream of the terminator hairpin and 3′-poly(U) tail. The secondary structures were predicted using *RNAstructure* 6.4 software and visualized using its Structure Editor (Reuter and Mathews 2010).

#### SUPPLEMENTAL TABLES

**Supplemental Table S1**. DNA oligonucleotides used to prepare templates for *in vitro* transcription.

| **Name** | **Sequence (5' -> 3')** |
|---|---|
| cspE81_wt_F | TAATACGACTCACTATAGGCCCTTCTGCTGCAAACGTAATCGCTCTGTAAGATACGTCAGCAAG |
| cspE81_wt_R | AAAAAAAACCCGCTGATTAAGCGGGTTTTGAATTCTTGCTGACGTATCTTACAGAG |
| cspE81-FinP-R | AAAATCCCCGCTGATTAAGCGGGTGTTGAATTCTTGCTGACGTATCTTACAGAG |
| cspE81-FinP-stem-R | AAAATCGCCGCTGATTAAGCGACTGTTGAATTCTTGCTGACGTATCTTACAGAG |
| FinP_wt_F | TAATACGACTCACTATAGGATACATAGGAACCTCCTCACAAAGGATTCTATGGACAGTCGATGCAGGG |
| FinP_wt_R | AAAATCGCCGATGCAGGGAGACGTGAACTCCCTGCATCGACTGTCCATAGAATCCT |
| FinP_malM_F | TAATACGACTCACTATAGGATACATAGGAACCTCCTCACAAAGGATTCTATGGACTGTCGATGCAGGG |
| FinP_malM_R | AAAAAAAGCCGATGCAGGGAGACGTGAACTCCCTGCATCGACAGTCCATAGAATCCT |
| FinP-U-UAU$_4$-R | AAAATAGCCGATGCAGGGAGACGTGAACTCCCTGCATCGACAGTCCATAGAATCCT |
| FinP-U-U$_6$-R | AAAAAAGCCGATGCAGGGAGACGTGAACTCCCTGCATCGACAGTCCATAGAATCCT |
| FinP-U-U$_4$- R | AAAAGCCGATGCAGGGAGACGTGAACTCCCTGCATCGACAGTCCATAGAATCCT |
| malM_wt_F | TAATACGACTCACTATAGCTTTATCAGCAGTGTAAAAGGCAAGGGGTAATTACGCCCCACAGTGCTGATT |
| malM_wt_R | AAAAAAAGGTGCGCCAGGAGACGCACCAGTTGTTGCAAAATCAGCACTGTGGGGCGTA |
| malM_FinP_R | AAAATCGGTGCGCCAGGAGACGCACCTGTTGTTGCAAAATCAG |

| | |
|---|---|
| | CACTGTGGGGCGTA |
| malM_RepX_R | AGAGCGGTGCGCCAGGAGACGCACCTAATGTTGCAAAATCAG CACTGTGGGGCGTA |
| malM-A-GU$_6$-R | AAAAAACGGTGCGCCAGGAGACGCACCTGTTGTTGCAAAATC AGCACTGTGGGGCGTA |
| malM-A-GAU$_5$-R | AAAAATCGGTGCGCCAGGAGACGCACCTGTTGTTGCAAAATCA GCACTGTGGGGCGTA |
| malM-A-GAU$_7$-R | AAAAAAATCGGTGCGCCAGGAGACGCACCTGTTGTTGCAAAAT CAGCACTGTGGGGCGTA |
| RepX_F | TAATACGACTCACTATAGCATTGATTGCCTCCTTTGCAGGCAGT TGGTGGTTAGGCGCTGGCGGGG |
| RepX_R | AGAGCGGCGCAGGGCGGGGGTAGTGACCCCGCCAGCGCCTAA CCACCAACTGCC |
| RepX-malM-F | TAATACGACTCACTATAGCATTGATTGCCTCCTTTGCAGGCAGT TGGTGGACTGGCGCTGGCGGGG |
| RepX-malM-R | AAAAAAAGGCGCAGGGCGGGGGTAGTGACCCCGCCAGCGCCA GTCCACCAAC |

**Supplemental Table S2.** DNA oligonucleotides used to prepare the insert to clone *E.coli fino* gene into the pET-15b expression plasmid.

| Name | Sequence (5' → 3') | Additional information |
|---|---|---|
| FinO-1_F | CTTTATTTCCAATCCATGACAGAGCAGAAGC | Insertion of a part of TEV Protease recognition sequence at the 5'-end of the coding sequence |
| FinO-2_F | GCCATATGCTCGA**GGATCC**gGAAAATCTTTATTTCCAA | Insertion of a part of TEV Protease recognition sequence and BamHI restriction site at the 5'-end of the coding sequence |
| FinO-1_R | GCATTGGTTCAAT**GGATCC**TTATTGCTCATCAAGCA | Insertion of the BamHI restriction site at the 3'-end of the coding sequence |

GAAAATCTTTATTTCCAATCC – TEV Protease recognition sequence

g – nucleotide introduced to maintain the open reading frame

**GGATCC** – BamHI restriction site

XXX – FinO coding sequence

XXX – additional sequences at 5'- and 3'-ends of the construct with FinO coding sequence inserted to improve the efficiency of BamHI cleavage

### SUPPLEMENTAL FIGURES

| Gene | Terminator hairpin | Gene | Terminator hairpin |
|---|---|---|---|
| cspC | 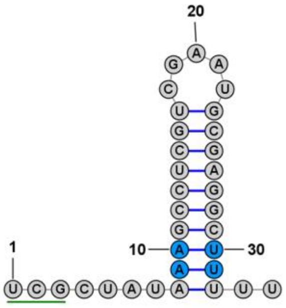 | cspE | 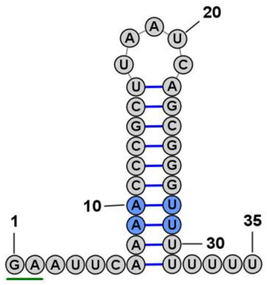 |
| uspA | 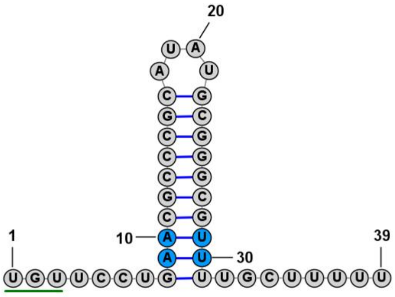 | rmf | 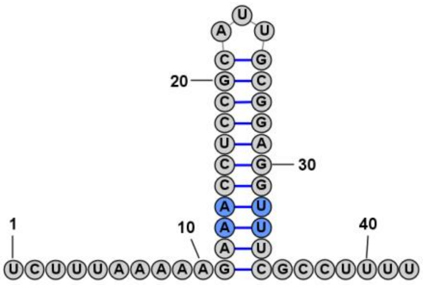 |
| csrB | 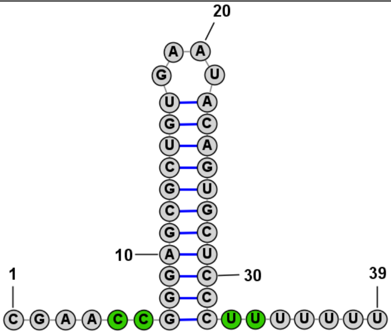 | ompC | 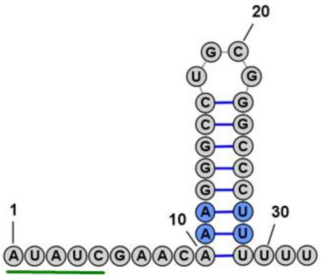 |
| gadC | 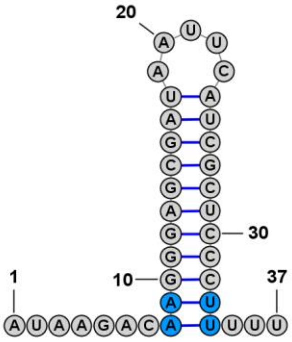 | dps | 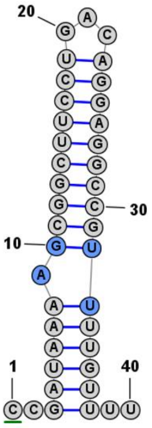 |

| | | | |
|---|---|---|---|
| **tnaA** | 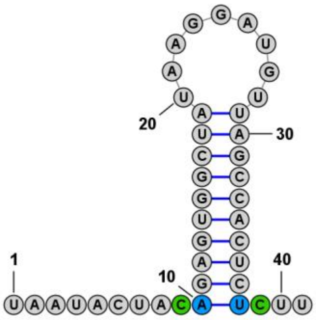 | **malM** | 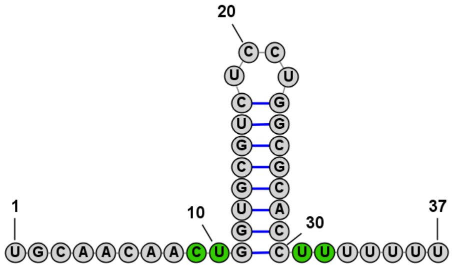 |
| **cspD** | 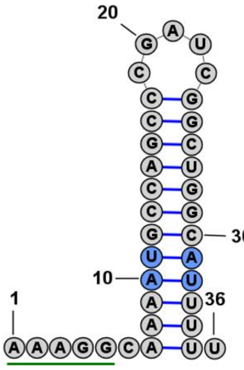 | **upd** | 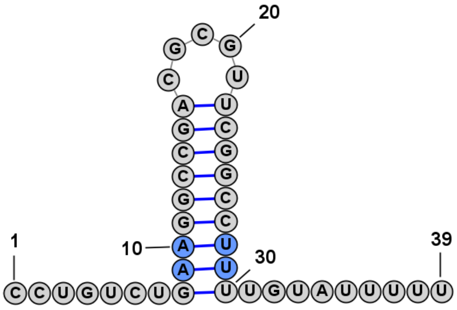 |
| **sibC** | 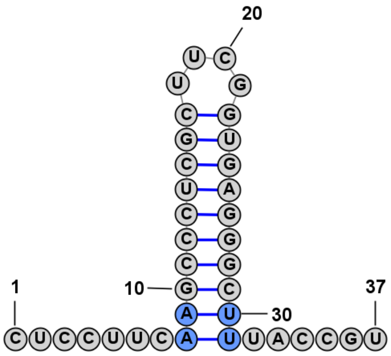 | **rpsQ** | 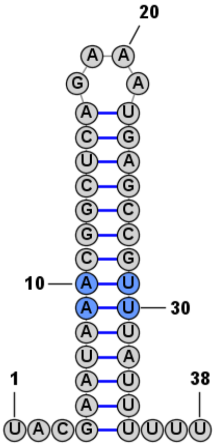 |
| **lpp** | 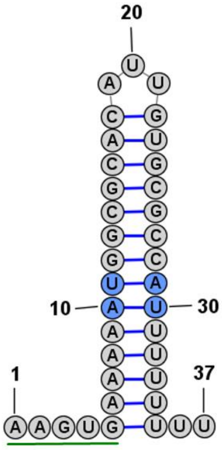 | **hupB** | 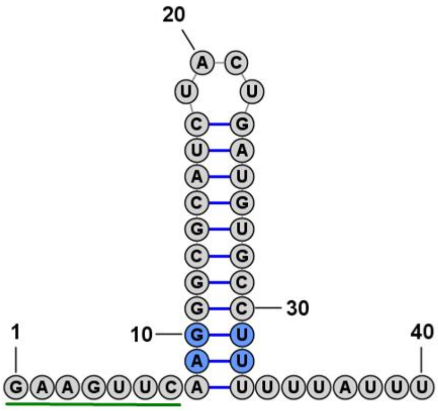 |

| | | | |
|---|---|---|---|
| **manZ** | 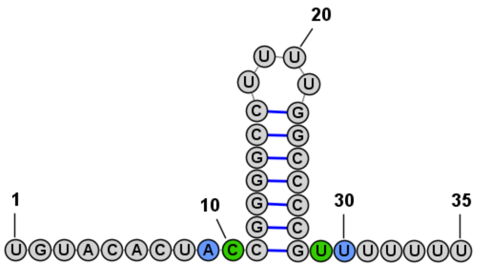 | **talB** | 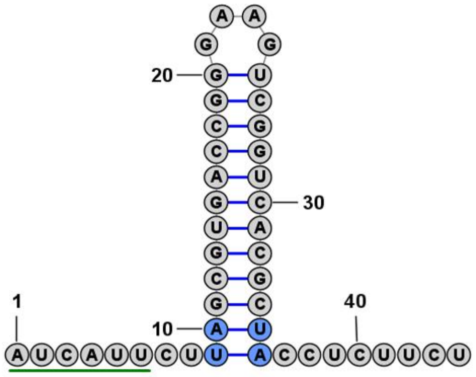 |
| **mcaS** | 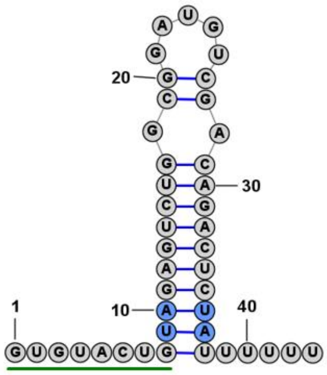 | **csrC** | 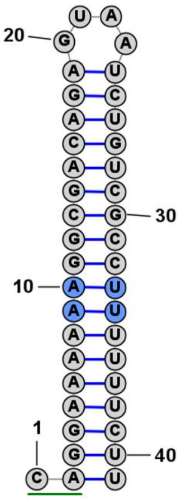 |

**Supplemental Figure S1.** Secondary structures of the terminator hairpins of the top 20 ProQ-specific RNA ligands identified in the previous CLIP-seq study in *Escherichia coli* (Holmqvist et al. 2018), whose peaks  map to the regions containing intrinsic transcription terminator. RNAs were ranked according to averaged read counts (Holmqvist et al. 2018). The secondary structures were predicted using the *RNAStructure* program (Reuter and Mathews 2010) for the 3′-terminal sequences, which included the terminator hairpin, the 3′ terminal polyU tail and a 10-nucleotide sequence upstream of the closing G-C or C-G base pair of the terminator hairpin. The two nucleotide pairs immediately below the closing G-C or C-G base pair of the terminator hairpin are marked in color, where blue denotes canonical base pairs (A-U, U-A, or G-U), and green denotes pyrimidine-pyrimidine appositions (U-U, C-U, or C-C). 5′ adjacent sequences, which are involved in secondary structure in the context of longer RNA are underlined green.

| Gene | Terminator hairpin | Gene | Terminator hairpin |
|---|---|---|---|
| **malM** | 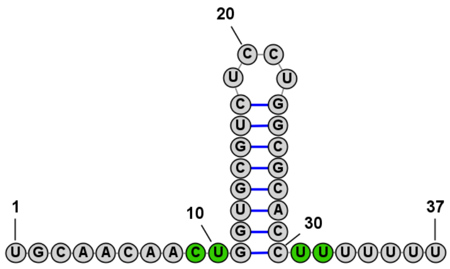 | **cspD** | 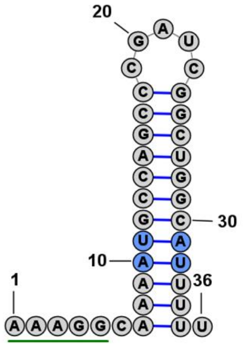 |
| **acpP** | 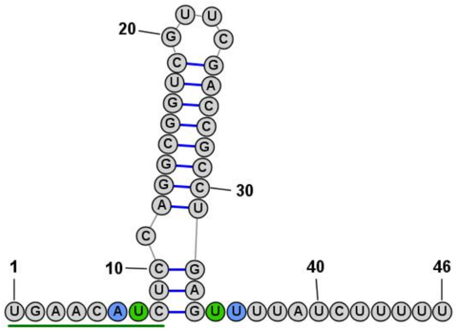 | **garD** | 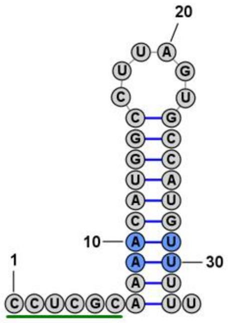 |
| **ryfA** | 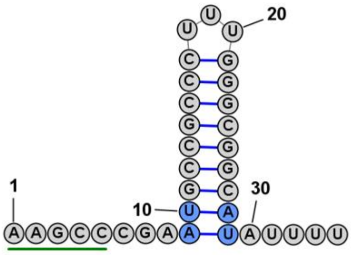 | **lpp** | 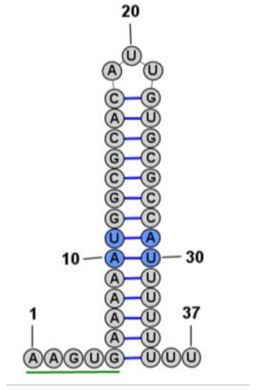 |
| **cspE** | 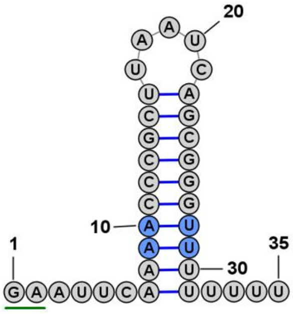 | **manZ** | 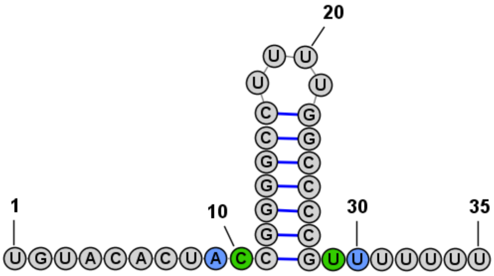 |

| | | | |
|---|---|---|---|
| **cspA** | 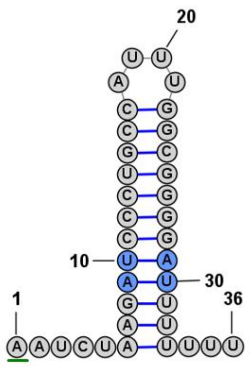 | **ompF** | 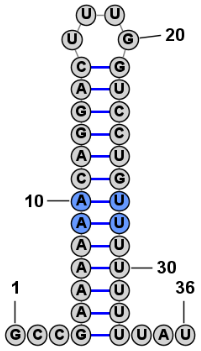 |
| **rybB** | 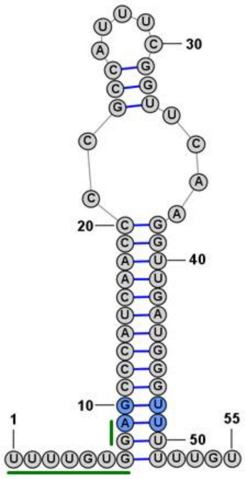 | **fbaA** | 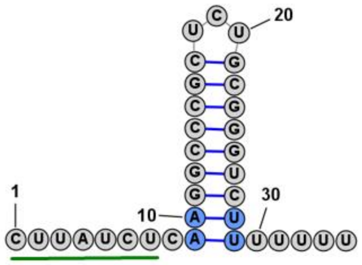 |
| **raiZ** | 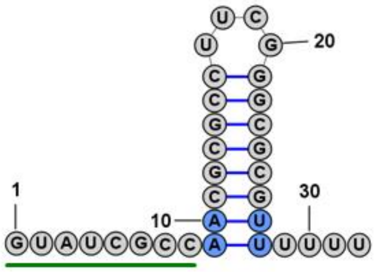 | **rraB** | 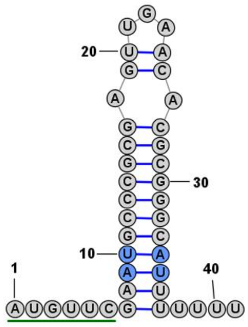 |
| **hupB** | 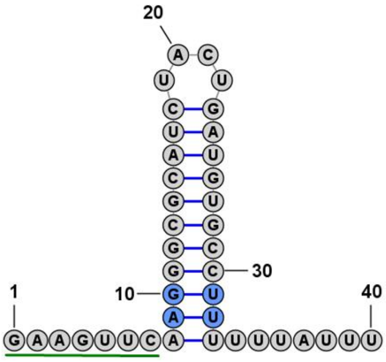 | **mcaS** | 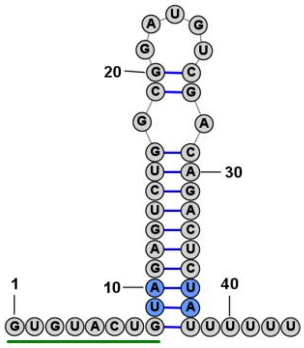 |

| | | | |
|---|---|---|---|
| **infA** | 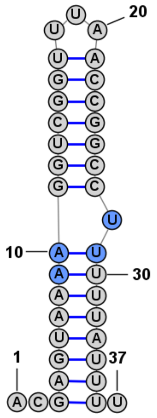 | **adhE** | 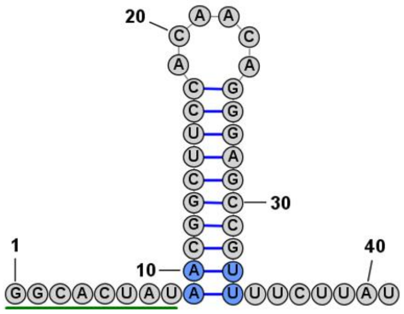 |
| **ihfA** | 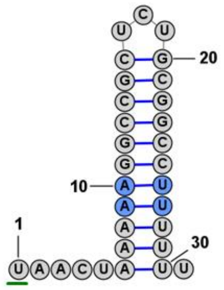 | **sodB** | 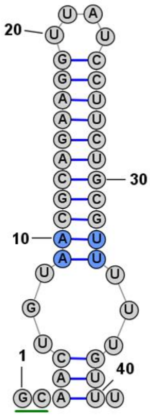 |

**Supplemental Figure S2.** Secondary structures of the terminator hairpins of the top 20 ProQ-specific RNA ligands identified in the previous RIL-seq study in *Escherichia coli* (Melamed et al. 2020), which were annotated either as 3′-UTRs or sRNAs. RNAs were ranked according to averaged read counts (Melamed et al. 2020). The secondary structures were predicted using the *RNAStructure* program (Reuter and Mathews 2010) for the 3′-terminal sequences, which included the terminator hairpin, the 3′ terminal polyU tail and a 10-nucleotide sequence upstream of the closing G-C or C-G base pair of the terminator hairpin. The two nucleotide pairs immediately below the closing G-C or C-G base pair of the terminator hairpin are marked in color, where blue denotes canonical base pairs (A-U, U-A, or G-U), and green denotes pyrimidine-pyrimidine appositions (U-U, or C-U). 5′ adjacent sequences, which are involved in secondary structure in the context of longer RNA are underlined green.

| sRNA | Terminator hairpin |
| --- | --- |
| **FinP** | 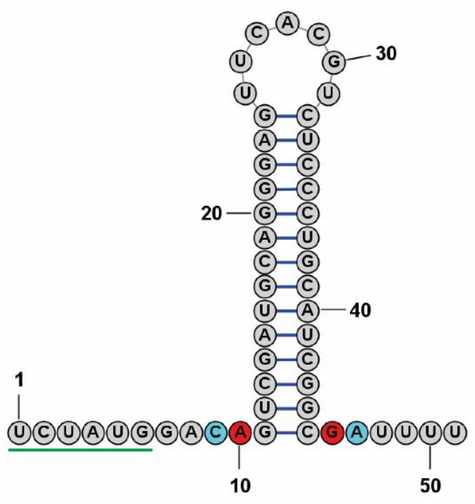 |
| **RepX** | 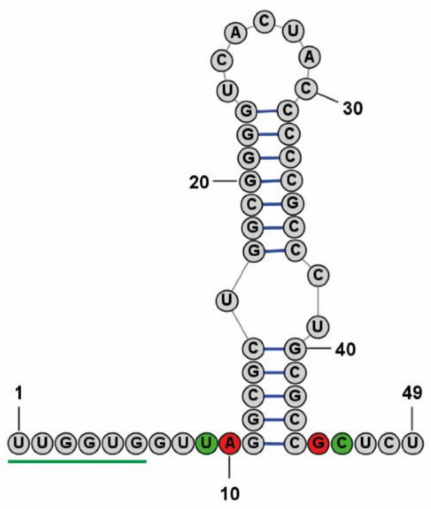 |

**Supplemental Figure S3.** Secondary structures of the two main *Salmonella enterica* FinO-specific RNA ligands, FinP and RepX, which were detected using RIP-seq (El Mouali et al. 2021). The secondary structures were predicted using the *RNAStructure* program (Reuter and Mathews 2010) for the 3′-terminal sequences, which included the terminator hairpin, the 3′ terminal tail and a 10-nucleotide sequence upstream of the closing G-C or C-G base pair of the terminator hairpin. The two nucleotide pairs immediately below the closing G-C or C-G base pair of the terminator hairpin are marked in color, where red denotes purine-purine appositions (A-G), green denotes a pyrimidine-pyrimidine apposition (U-C), and light blue denotes a C-A apposition. 5′ adjacent sequences, which are involved in secondary structure in the context of longer RNA are underlined green.

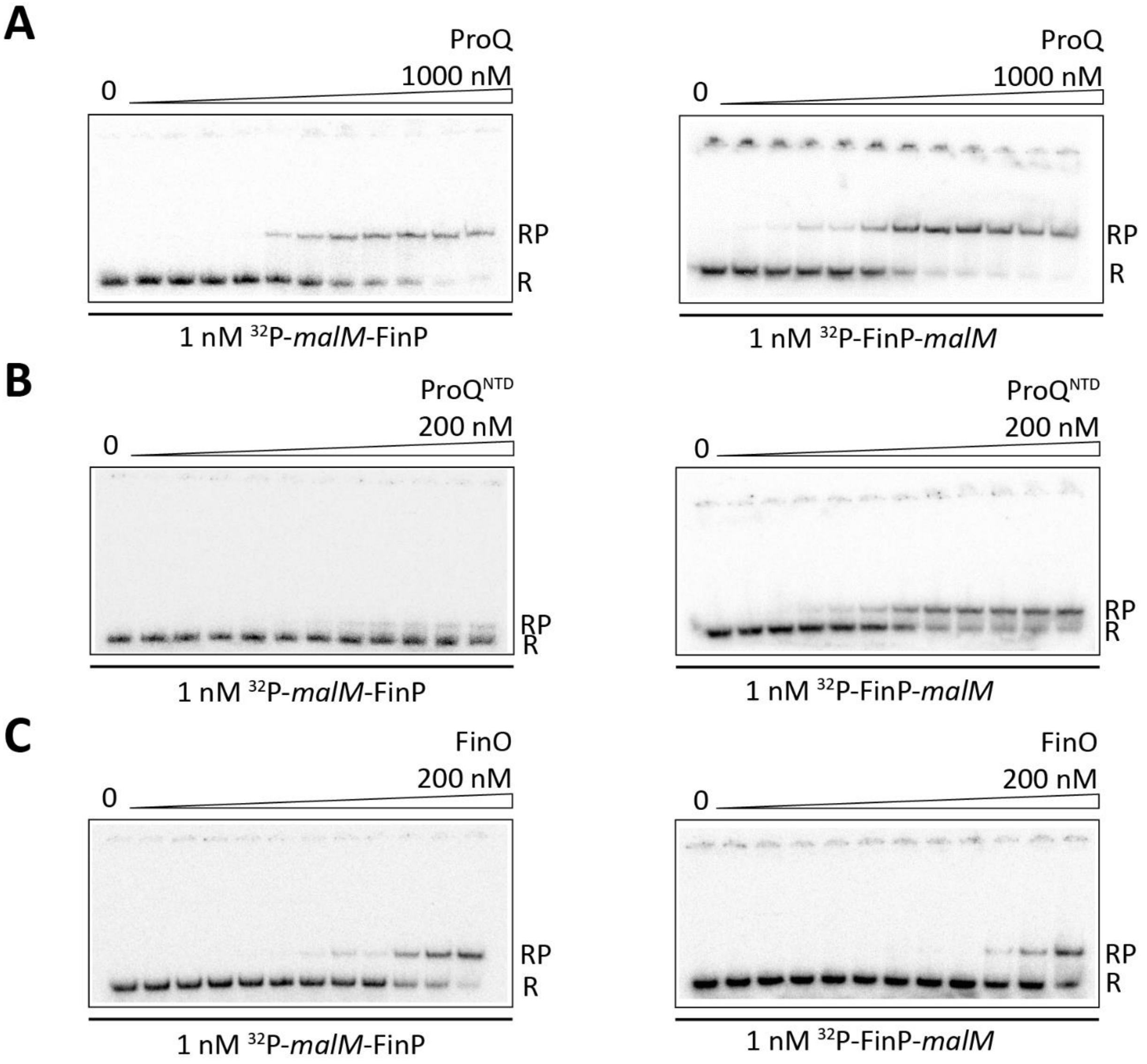


**Supplemental Figure S4.** Gelshift analysis of the binding of 1 nM *malM*-FinP and FinP-*malM* binding to full-length ProQ (A), ProQ$^{NTD}$ (B), and FinO (C). Free $^{32}$P-labeled RNA is marked as R and RNA-protein complexes as RP. The fitting of *malM*-FinP data using the quadratic equation provided $K_d$ value of 85 nM for binding to ProQ and 83 nM for binding to FinO, while the $K_d$ value for binding to ProQ$^{NTD}$ was estimated as higher than 200 nM. The fitting of FinP-*malM* data using the quadratic equation provided $K_d$ value of 27 nM for binding to ProQ, 4.9 nM for binding to ProQ$^{NTD}$, and 327 nM for binding to FinO. The average equilibrium dissociation constant ($K_d$) values calculated from at least three independent experiments are shown in Table 1 in the main text.

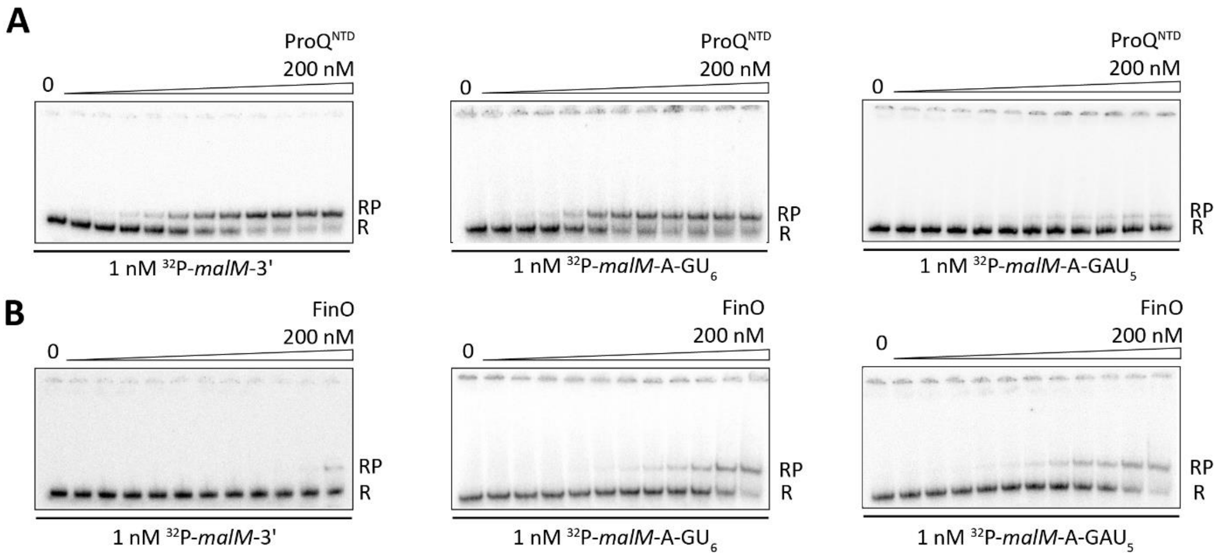


**Supplemental Figure S5.** Gelshift analysis of the binding of 1 nM *malM*-3', *malM*-A-$GU_6$ and *malM*-A-$GAU_5$ binding to $ProQ^{NTD}$ (A), and FinO (B). Free $^{32}$P-labeled RNA is marked as R and RNA-protein complexes as RP. The fitting of *malM*-A-$GU_6$ data using the quadratic equation provided $K_d$ value of 1.8 nM for binding to $ProQ^{NTD}$, and 151 nM for binding to FinO. The fitting of *malM*-A-$GAU_5$ data using the quadratic equation provided $K_d$ value of 122 nM for binding to FinO, while the $K_d$ value for binding to $ProQ^{NTD}$ was estimated higher than 200 nM. The data shown for *malM*-3' are the same as in Figure 1. The average equilibrium dissociation constant ($K_d$) values calculated from at least three independent experiments are shown in Table 1 in the main text.

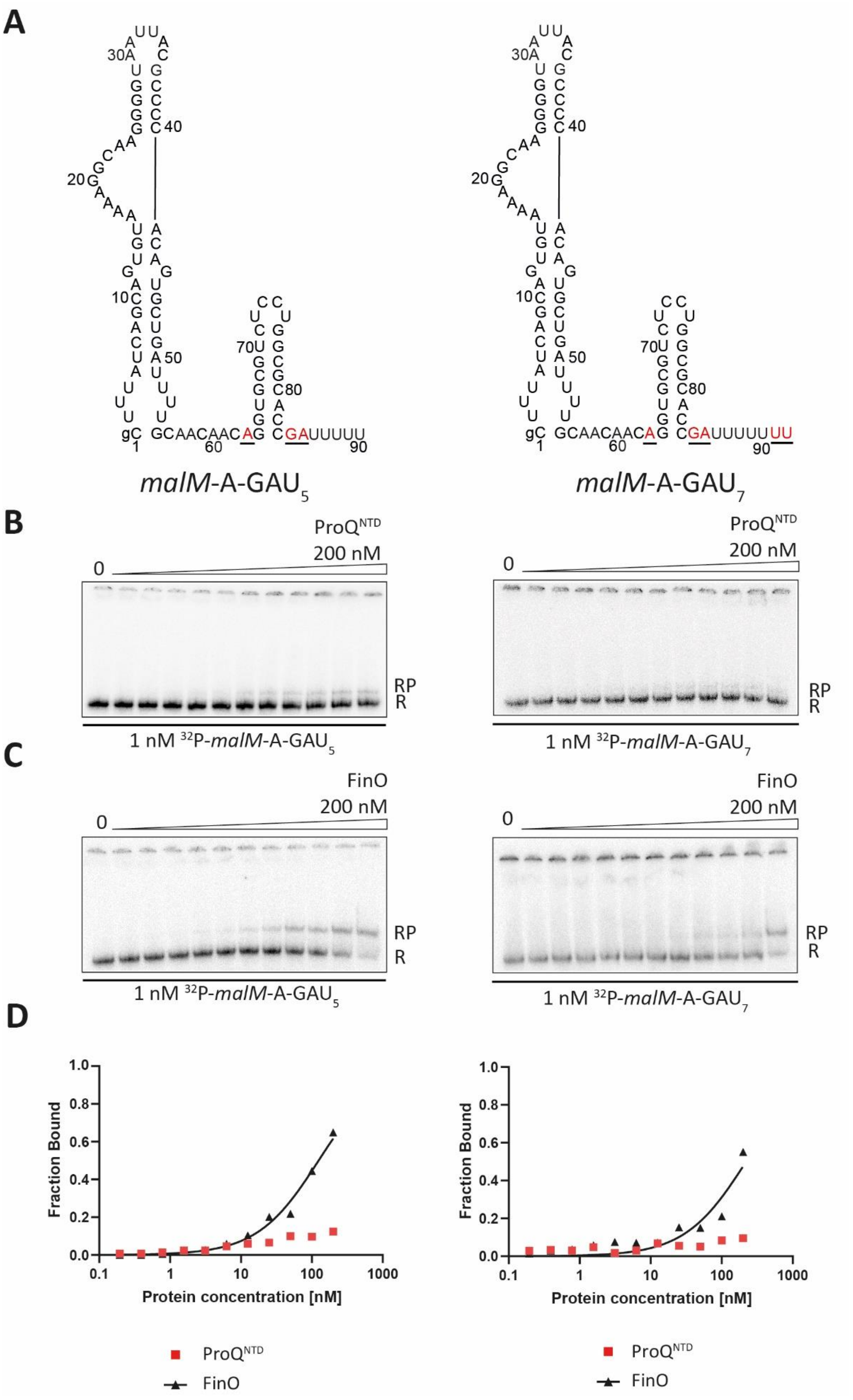


**Supplemental Figure S6.** Comparison of *malM*-A-GAU$_5$ and *malM*-A-GAU$_7$ binding to ProQ$^{NTD}$, and to FinO. (A) Secondary structures of *malM*-A-GAU$_5$ and *malM*-A-GAU$_7$, which were predicted using *RNAstructure* software (Reuter and Mathews 2010). The substitutions introduced into *malM*-3' are shown in red, underlined font. The lower case g denotes guanosine residue added on 5' ends of RNA molecules to enable T7 RNA polymerase transcription. (B), (C) The gelshift analysis of *malM*-A-$GAU_5$ and *malM*-A-$GAU_7$ binding to $ProQ^{NTD}$ (B), and FinO (C). (D) The plots of fraction bound data from (B) and (C) versus protein concentration are shown for *malM*-A-$GAU_5$ and *malM*-A-$GAU_7$. The data shown for *malM*-A-$GAU_5$ are the same as in Figure 3 (main text), and the corresponding average $K_d$ values are shown in Table 1 (main text). Because the maximum fraction bound of *malM*-$GAU_7$ binding to $ProQ^{NTD}$ was 8%, the $K_d$ value was assumed as bigger than 200 nM. The data for *malM*-A-$GAU_7$ binding to FinO were analyzed by fitting to the quadratic equation, which provided the $K_d$ value of 197 ± 86 with maximum fraction bound of 56%. The values were calculated from at least three independent experiments.

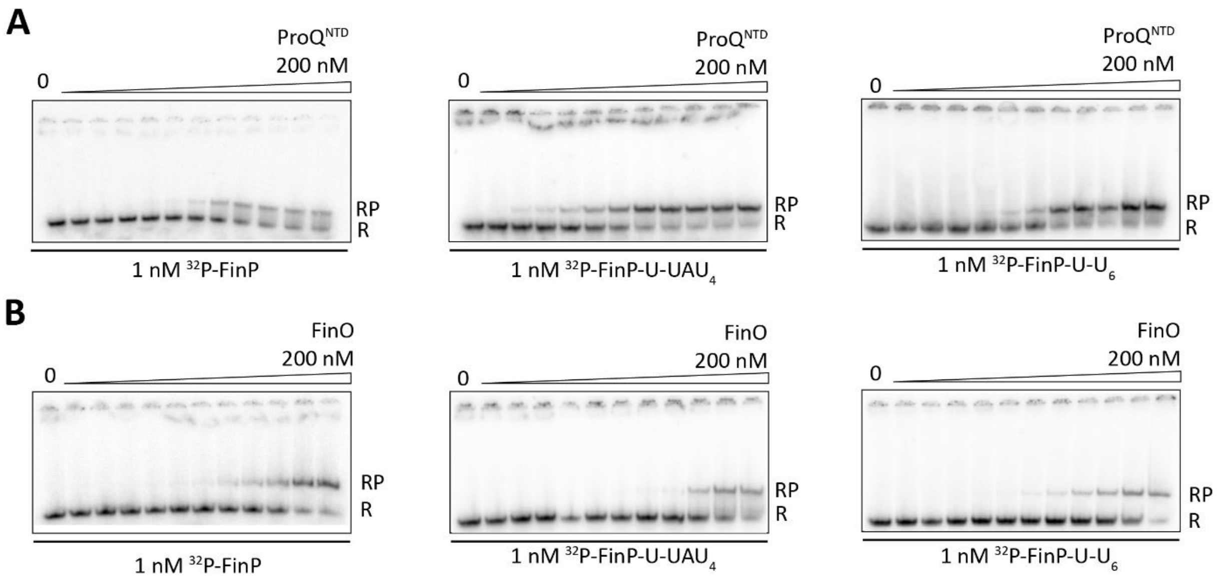


**Supplemental Figure S7.** Gelshift analysis of the binding of 1 nM FinP, FinP-U-UAU$_4$ and FinP-U-U$_6$ binding to ProQ$^{NTD}$ (A), and FinO (B). Free $^{32}$P-labeled RNA is marked as R and RNA-protein complexes as RP. The fitting of FinP-U-UAU$_4$ data using the quadratic equation provided $K_d$ value of 4.6 nM for binding to ProQ$^{NTD}$, and 184 nM for binding to FinO. The fitting of FinP-U-U$_6$ data using the quadratic equation provided $K_d$ value of 11 nM for binding to ProQ$^{NTD}$, and 160 nM for binding to FinO. The data shown for FinP are the same as in Figure 1. The average equilibrium dissociation constant ($K_d$) values calculated from at least three independent experiments are shown in Table 1 in the main text.

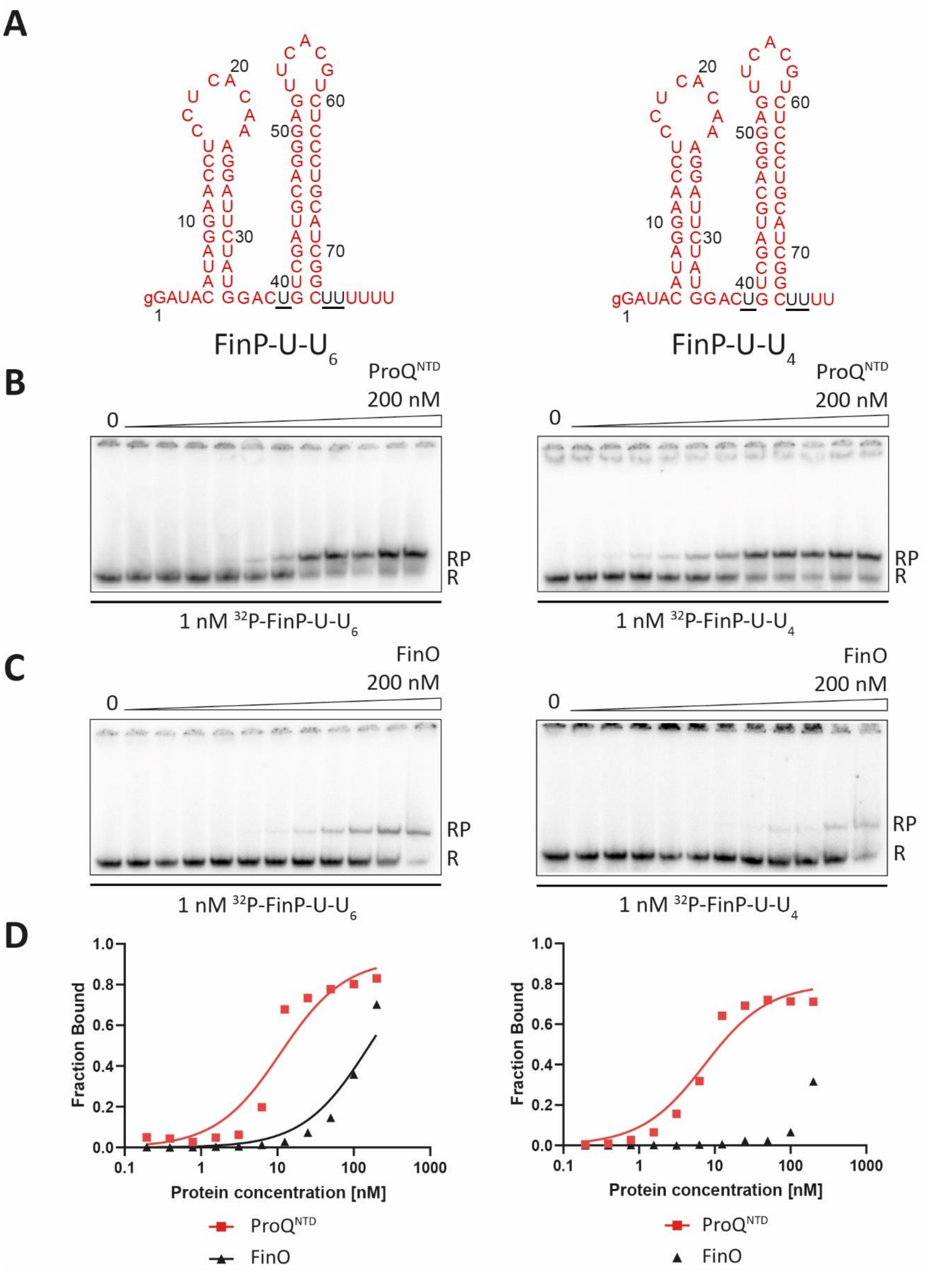


**Supplemental Figure S8.** Comparison of FinP-U-U$_6$ and FinP-U-U$_4$ binding to ProQ$^{NTD}$ and to FinO. (A) Secondary structures of FinP-U-U$_6$ and FinP-U-U$_4$, which were predicted using *RNAstructure* software (Reuter and Mathews 2010). The substitutions introduced into FinP are shown in black, underlined font. The lower case g denotes guanosine residue added on 5' ends of RNA molecules to enable T7 RNA polymerase transcription. (B), (C) The gelshift analysis of FinP-U-U$_6$ and FinP-U-U$_4$ binding to ProQ$^{NTD}$ (B), and FinO (C). (D) The plots of fraction bound data from (B) and (C) versus protein concentration are shown for FinP-U-$U_6$ and FinP-U-$U_4$. The data shown for FinP-U-$U_6$ are the same as in Figure 3 (main text), and the corresponding average $K_d$ values are shown in Table 1 (main text). The data for FinP-U-$U_4$ binding to ProQ$^{NTD}$ were analyzed by fitting to the quadratic equation, which provided the $K_d$ value of 7.6 ± 3.0 nM with maximum fraction bound of 71%. Because the maximum fraction bound of FinP-U-$U_4$ binding to FinO was 30%, the $K_d$ value was assumed as bigger than 200 nM. The $K_d$ values were calculated from at least three independent experiments.

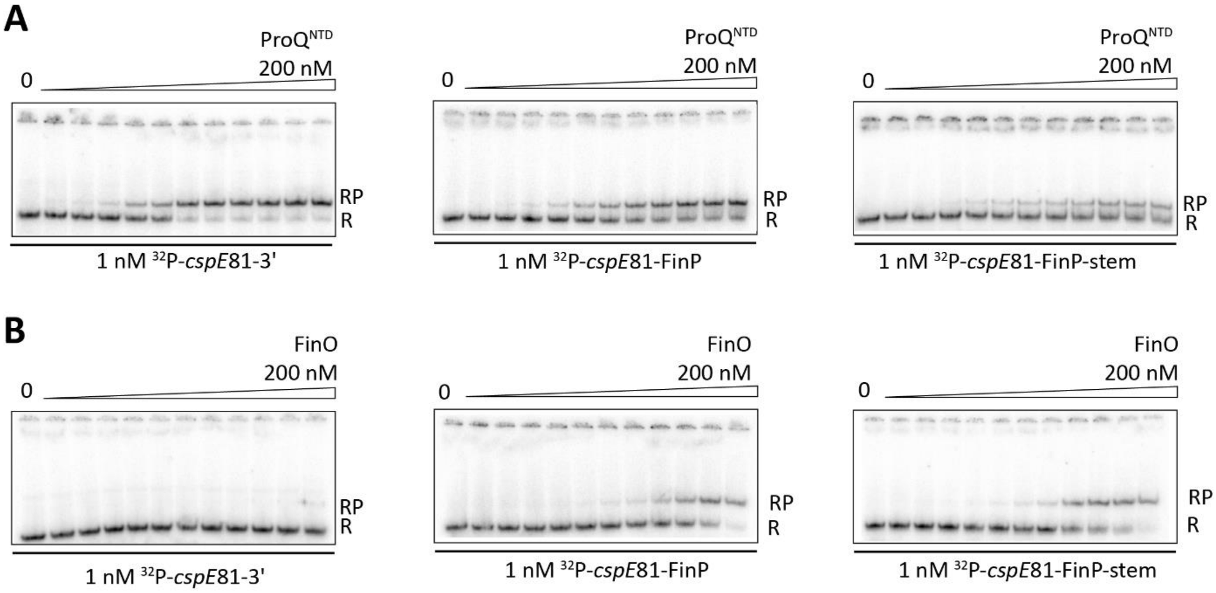


**Supplemental Figure S9.** Gelshift analysis of the binding of 1 nM *cspE*81-3', *cspE*81-FinP and *cspE*81-FinP-stem binding to ProQ$^{NTD}$ (A), and FinO (B). Free $^{32}$P-labeled RNA is marked as R and RNA-protein complexes as RP. The fitting of *cspE*81-3' data using the quadratic equation provided $K_d$ value of 2.7 nM for binding to ProQ$^{NTD}$, while the $K_d$ value for binding to FinO was estimated as higher than 200 nM. The fitting of *cspE*81-FinP data using the quadratic equation provided $K_d$ value of 5.0 nM for binding to ProQ$^{NTD}$, and 99 nM for binding to FinO. The fitting of *cspE*81-FinP-stem data using the quadratic equation provided $K_d$ value of 55 nM for binding to FinO, while the $K_d$ value for binding to ProQ$^{NTD}$ was estimated as higher than 200 nM. The average equilibrium dissociation constant ($K_d$) values calculated from at least three independent experiments are shown in Table 1 in the main text.

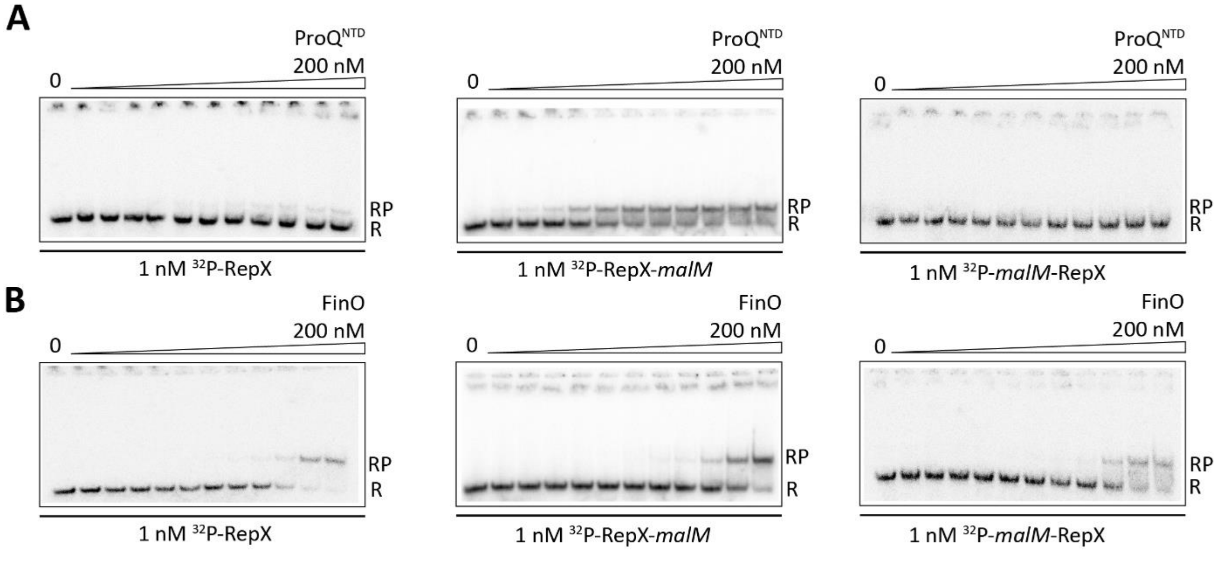


**Supplemental Figure S10.** Gelshift analysis of the binding of 1 nM RepX, RepX-*malM* and *malM*-RepX binding to ProQ$^{NTD}$ (A), and FinO (B). Free $^{32}$P-labeled RNA is marked as R and RNA-protein complexes as RP. The fitting of RepX data using the quadratic equation provided $K_d$ value of 79 nM for binding to FinO, while the $K_d$ value for binding to ProQ$^{NTD}$ was estimated as higher than 200 nM. The fitting of RepX-*malM* data using the quadratic equation provided $K_d$ value of 3.0 nM for binding to ProQ$^{NTD}$, and 148 nM for binding to FinO. The fitting of *malM*-RepX data using the quadratic equation provided $K_d$ value of 137 nM for binding to FinO, while the $K_d$ value for binding to ProQ$^{NTD}$ was estimated as higher than 200 nM. The average equilibrium dissociation constant ($K_d$) values calculated from at least three independent experiments are shown in Table 1 in the main text.

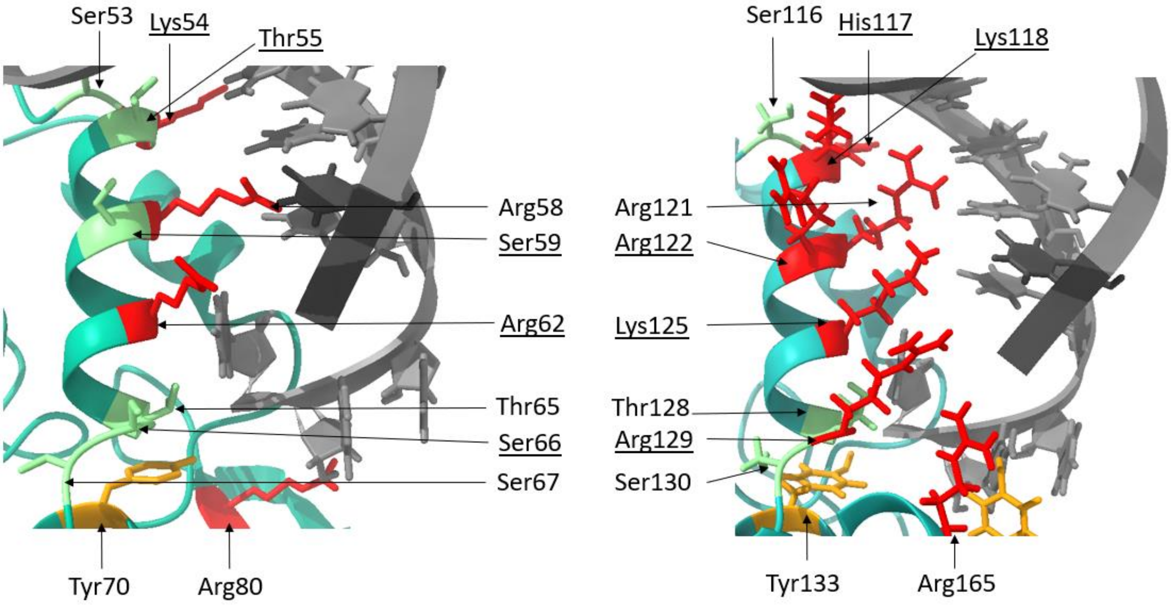


**Supplemental Figure S11.** The modeling of RNA binding surfaces in ColabFold-predicted structures of *E. coli* ProQ and F-like plasmid FinO. The figure shows the α-helix 3 from ProQ (left) and the corresponding α-helix 4 from FinO (right) with marked amino acid residues pointing towards modeled location of RNA helix. Both structures were predicted using ColabFold software (Mirdita et al. 2022). The modelling of interactions was done using Chimera X (Pettersen et al. 2021) by aligning the ColabFold-predicted structure of the FinO domain of *E. coli* ProQ (A) and the ColabFold-predicted structure of the F-like plasmid FinO protein (B) with the X-ray structure of the FinO domain of *L. pneumophila* RocC in complex with the terminator hairpin of RocR RNA (Kim et al. 2022). The side chains of amino acid residues located in the corresponding positions of both proteins are marked in color, with arginine, lysine and histidine residues marked in red, serine and threonine in green, and tyrosine in orange. The descriptions of corresponding amino acids are located in corresponding places on the figure. The descriptions of those amino acids, which are located in corresponding positions, but are different, are underlined. The structure of the *L. pneumophila* RocR hairpin is shown in grey.
